# Unveiling *Cryptosporidium parvum* Sporozoite-Derived Extracellular Vesicles: Profiling, Origin, and Protein Composition

**DOI:** 10.1101/2023.12.21.571703

**Authors:** Lucia Bertuccini, Zaira Boussadia, Anna Maria Salzano, Ilaria Vanni, Ilaria Passerò, Emanuela Nocita, Andrea Scaloni, Massimo Sanchez, Massimo Sargiacomo, Maria Luisa Fiani, Fabio Tosini

## Abstract

*Cryptosporidium parvum* is a common cause of a zoonotic disease and a main cause of diarrhea in newborns around the world. Effective drugs or vaccines are still lacking. Oocyst is the infective form of the parasite; after the ingestion the oocyst excysts and releases four sporozoites into the intestine that rapidly attack the enterocytes. The membrane protein CpRom1 is a large rhomboid protease that is expressed by sporozoites and recognized as antigen by the host immune system. In this study, we observed the release of CpRom1 with extracellular vesicles (EVs) not previously described. To investigate this phenomenon, we isolated and resolved EVs from the excystation medium by differential ultracentrifugation. By fluorescence flow cytometry and transmission electron microscopy (TEM), we identified two types of sporozoite-derived vesicles: large extracellular vesicles (LEVs) and small extracellular vesicles (SEVs), having different a mean size of 150 nm and 60 nm, respectively. Immunodetection experiments proved the occurrence of CpRom1 and the Golgi protein CpGRASP in LEVs, while immune-electron microscopy experiments demonstrated localization of CpRom1 on LEVs surface.

TEM and scanning electron microscopy (SEM) showed the generation of LEVs by budding of the outer membrane of sporozoites; conversely, the origin of SEVs remained uncertain.

Differences between LEVs and SEVs were observed for protein composition as proved by the corresponding electrophoretic profiles. Indeed, a dedicated proteomic analysis identified 5 proteins unique to LEVs composition and 16 proteins unique to SEVs. Overall, 60 proteins were identified in the proteome of both types of vesicles and most of these proteins (48 in number) were already identified in the molecular cargo of extracellular vesicles from other organisms. Noteworthy, we identified 12 proteins unique to *Cryptosporidium* spp. that had never been associated with EVs. This last group included the immunodominant parasite antigen glycoprotein GP60, which is one of the most abundant proteins in both LEVs and SEVs.

## 1 Introduction

*Cryptosporidium* genus consists of 38 species of obligate parasites of the phylum Apicomplexa that infect vertebrates of many species (Feng et al., 2018). Several species of *Cryptosporidium* are important pathogens affecting mammals’ intestine and cause cryptosporidiosis, a diarrhoeal disease that usually lasts several days. Two closely related species, namely *Cryptosporidium parvum* and *Cryptosporidium hominis*, are responsible for most cases of human cryptosporidiosis with watery diarrhoea as the main symptom (Bouzid et al., 2013). In immunocompetent adults, cryptosporidiosis usually has a limited duration (7-10 days), but differently, this infection poses a serious risk to people with immunodeficiency, who may have a long-term and life-threatening illness due to severe dehydration (Checkley et al., 2015). Remarkably, *C. parvum* and *C. hominis* are also responsible for most cases of neonatal cryptosporidiosis, a major cause of diarrhoea in children in endemic regions, where transmission is favoured by poor hygiene conditions (Kotloff et al., 2013). Transmission of the parasite occurs via the oral-fecal route either directly (animal to human or human to human) or indirectly through contaminated food and, above all, water (Xiao and Feng, 2017).

Oocyst is the parasitic stage released with feces by infected subjects, and it is responsible for the transmission of the infection to other subjects. Oocysts are characterized by a thick “shell”, the oocyst wall, constituted by polysaccharides, lipids and peculiar proteins defined as *Cryptosporidium* oocyst wall proteins (COWPs), which make oocysts particularly resistant to environmental and chemical stresses (Templeton et al., 2004). Noteworthy, oocysts are also resistant to treatments to disinfect drinkable water; this is a significant risk factor for water plant contamination, which can cause large outbreaks with thousands of infected people (MacKenzie et al., 1994). Despite the relevance of *Cryptosporidium* for human health, effective drug therapies are still lacking, and vaccines have not yet been developed.

*Contrarily to other apicomplexan parasites such as Plasmodium spp. and Toxoplasma gondii, Cryptospordium* completes its life cycle in a single host (monoxenous). Excystation is the first stage of the infectious process and consists of rupture of the oocyst wall and the egress of four sporozoites; this is triggered first the passage of the oocyst in the stomach then by its subsequent contact with the bile salts in the small intestine (Fayer and Xiao, 2007). The released sporozoites adhere to the luminal side of enterocytes in the small intestine. The close contact between the parasite and the host cell membrane induces the formation of a parasitophorous vacuole (PV). PV protrudes toward the intestinal lumen, but the intracellular stages of the parasite are completely wrapped by the host cell membrane. The motile stages of *Cryptosporidium* (*i. e.*, sporozoites and merozoites), like other apicomplexans, are characterized by the apical complex at the anterior end of the cell. The apical complex includes specialized subcellular organelles, such as rhoptry, numerous micronemes and various dense granules, which are vesicular structures specialized for the parasitic function. These organelles discharge their content sequentially from the egress of sporozoites during the attachment and invasion of the host cell, and until the formation of a parasitophorous vacuole at the luminal side of the cell (Guerin et al., 2023).

Oocyst excystation as well as the organelle discharge can be easily replicated *in vitro* even in the absence of host cells. In fact, free sporozoites maintained at 37 °C in an adequate medium, release most of their organelle content within two hours (Chen et al., 2004). Free sporozoites, among other things, carry antigens that can be successfully counterbalanced by circulating host antibodies (Tosini et al., 2019). By investigating the antigenic proteins involved in the preliminary phases of the excystation, we identified a rhomboid protease, namely CpRom1, which occurs in sporozoites (Trasarti et al., 2007). Rhomboids are ubiquitous serine-proteases that are entirely embedded in the lipid bilayer of cell membranes (Kühnle et al., 2019). Rhomboids in Apicomplexa play a unique role before and during the invasion of the host cell by cleaving adhesins, surface proteins exposed in the motile stages such as sporozoites, during their path towards the host cell and in the phase of penetration through the host cell membrane (Carruthers and Blackman 2005; Dowse et al., 2008).

In this study, we observed that CpRom1 is released by sporozoites as bound to extracellular vesicles (EVs). This coincidence led us to investigate the release of microvesicles by sporozoites. Nowadays, EVs are recognized as fundamental elements for intercellular communication both in unicellular and in multicellular organisms (Doyle and Wang, 2019). They also serve as effectors of intercellular communication in protozoan pathogens, mediating interactions between parasites and between parasites and host cells (Wang et al., 2022). Here we describe that *C. parvum* sporozoites release two types of EVs that, based on their size, we have here classified as large extracellular vesicles (LEVs) and small extracellular vesicles (SEVs). These vesicles exhibited significant differences in their protein composition, but also presented common molecules including one of the most antigenic *C. parvum* component, namely glycoprotein GP60 (Allison et al., 2011), also known as GP40/GP15.

## 2 Materials and Methods

### 2.1 Parasites and excystation procedure

Fresh *C. parvum* oocysts (Iowa strain) were supplied by Bunch Grass Farm (Deary, Idaho USA), stored at 4 °C in PBS with penicillin (1000 U.I./ml) and streptomycin (1 mg/ml). The excystation procedure was performed as follows: aliquots of 1x10^7^ oocysts per ml were pelleted at 2000 rpm, for 5 min, resuspended in 1 ml of 10 mM HCl, and then incubated at 37 °C for 10 min. Oocysts were pelleted again as above and resuspended in 1 ml of excystation medium (D-MEM containing 2 mM sodium taurocholate); they were maintained at 15 °C, for 10 min, and then moved to 37 °C to induce the excystation. Excystation mixtures were sampled at various times after the induction.

### 2.2 Expression of recombinant CpRom1 and CpGRASP and production of the corresponding antisera

The CpRom1 and CpGRASP coding seqence (monoexonic genes without introns) were directly amplified from *C. parvum* genomic DNA (Iowa strain).

#### 2.2.1 Cloning, expression and purification of recombinant CpRom1

The recombinant 6His-CpRom1 was obtained as follows: the CpRom1 coding sequence was amplified with AAC<u>GAGCTC</u>GATATGTCCGATTTTGTTTTCA as the forward primer including the *Sac*I (underlined) restriction site, and TCCC<u>CCCGGG</u>TCATCAAGAAAAATCATATCCAAATA as the reverse primer including the *Sma*I (underlined) restriction site. Amplification with 2X Phusion Flash High fidelity PCR Master Mix (Finnzymes) was performed using 80 ng of genomic DNA as template as follows: 95 °C for 5 min, 35 cycles of 94 °C for 15 sec, 50 °C for 15 sec, 72 °C for 4 min, and a final extension at 72 °C for 10 min in a Veriti 96 well thermal cycler (Applied Biosystem). Amplicons were purified with QIAquick PCR Purification Kit (Qiagen GmbH, Hilden, Germany) and digested with *Sac*I and *Sma*I restriction enzymes (New England Biolabs). This fragment was then ligated in the *Sac*I and *Sma*I digested pQE80 vector (Qiagen GmbH, Hilden, Germany) using the Quick Ligase kit (New England Biolabs), and the ligation mix was used to transform the *Escherichia coli* M15 strain. Positive clones were selected on LB agar plates with 100 µg/ml ampicillin and 25 µg/ml kanamycin by PCR screening; recombinant plasmid with recombinant CpRom1 ORF was sequenced to check the fusion with the histidine-tag coding sequences at its N-terminus. For 6His-CpRom1 purification, 20 ml of an overnight culture of recombinant bacteria were inoculated in 1 l of LB with 100 µg/ml ampicillin and 25 µg/ml kanamycin, and cultured at 37 °C, with vigorous shacking, until a 0.6 OD was reached; then, 1 mM IPTG was added to the culture and the bacterial growth continued for additional 3 h. Bacteria were pelleted first at 4,000 rpm for 10 min, resuspended in 100 ml cold PBS and centrifuged again at 6,000 rpm for 20 min. Then, the pellet was resuspended in 20 ml of denaturing buffer A (100 mM NaH_2_PO4, 10 mM Tris-HCl, 6 M guanidine-HCl, pH 8.0), and stirred at 25 °C, overnight; the corresponding lysate was clarified by centrifugation at 10,000 rpm, for 30 min. Then, 5 ml of 50% Ni-NTA resin was added and the mix was gently stirred for 60 min, at 25°C; the slurry was loaded slowly on a 10 ml column to pack the resin and then washed with 40 ml (8 x 5 ml wash) of buffer C (100 mM NaH_2_PO4, 10 mM Tris-HCl, 8 M urea, pH 6.3). Purified protein was eluted with 4 aliquots of 2.5 ml of buffer D (100 mM NaH_2_PO4, 10 mM Tris-HCl, 8 M urea, pH 5.9) and with 4 aliquots of 2.5 ml of buffer E (100 mM NaH_2_PO4, 10 mM Tris-HCl, 8 M urea, pH 4.5). Pooled fractions from buffer D and pooled fractions from buffer E were extensively dialyzed against PBS with increasing concentrations of glycerol (from 10% to 50%) in Slide-A-Lizer^tm^ (ThermoFisher) dialysis cassettes with a cut-off of 10 kDa.

#### 2.2.2 Cloning, expression and purification of recombinant CpGRASP

To clone the CpGRASP coding sequence, the following forward primer CC<u>GGATCC</u>GGAGGTGCGCAAACCAAAC including *Bam*HI restriction site (underlined) and reverse primer CT<u>CCCGGG</u>TTATATTTCTCCTTGGTCTGTG including *Sma*I restriction site (underlined) were used. PCR amplification was performed with Hot Star Taq Plus (Qiagen) using 5 ng of genomic DNA as template and these PCR conditions: 95 °C for 5 min, 35 cycles of 94 °C for 15 sec, 55 °C for 15 sec, 72 °C for 4 min, and a final extension at 72 °C for 10 min. Reactions were conducted in a Veriti 96 well thermal cycler (Applied Biosystem). The amplified fragment was digested with *Bam*HI and *Sma*I restriction enzymes and ligated in pQE80 vector (Qiagen) digested with the same enzymes, using Quick Ligase kit (New England Biolabs). Ligation was used to transform *E. coli* M15 host strain, positive colonies were selected by PCR screening, and amplicons were sequenced to verify the fusion of histidine tag at the 5’-end of the inserted CpGRASP sequence. The histidine-tagged CpGRasp (6His-CpGRASP) was then purified as above.

#### 2.2.3 Production of specific mouse antisera for 6His-CpRom1 and 6His-CpGRASP

To produce specific antisera for 6His-CpRom1 and 6His-CpGRASP, Balb-C mice were immunized with the following schedule: i) 100 mg of protein plus complete Freund adjuvant as first inoculum; ii) 100 mg of protein plus incomplete Freund adjuvant as second inoculum, 30 days after the first inoculum; iii) a final inoculum with 100 mg of protein in PBS, 30 days after the second inoculum. Mice were bled 15 days after the last inoculum and 150-300 ml of serum was obtained from each mouse.

### 2.3 Isolation of extracellular vesicles from excystation medium by differential centrifugation

Sporozoite microvesicles were prepared from the excystation medium of fresh oocyst aliquots after 2 h incubation at 37 °C; excystation was blocked by placing samples on ice for 5 min. All subsequent manipulations were performed on ice and centrifugation steps were performed at 4 °C. Excystation medium was centrifuged at 2000 rpm, for 20 min, to remove most of oocysts and sporozoites and the resulting supernatant was centrifuged again at 5000 rpm, for 10 min, to remove residual oocysts, sporozoites, and excystation debris. To isolate EVs, supernatant was transferred to 1.5 ml tubes (Eppendorf), centrifuged for 20 min at 10,000 g and the pellet containing LEVs resuspended in 100 μl of ice-cold PBS. To isolate SEVs, the supernatant was transferred to new tubes (Beckman Coulter) and ultracentrifuged on a TLA 120.2 rotor (Beckman Coulter) at 100,000 g for 3 hours. The pellet was resuspended in 1.2 ml of ice-cold PBS and recentrifuged as above. The final pellet containing SEVs was resuspended in 100 μl of ice-cold PBS. All the procedure is schematized in Figure 1.

**Figure 1.**
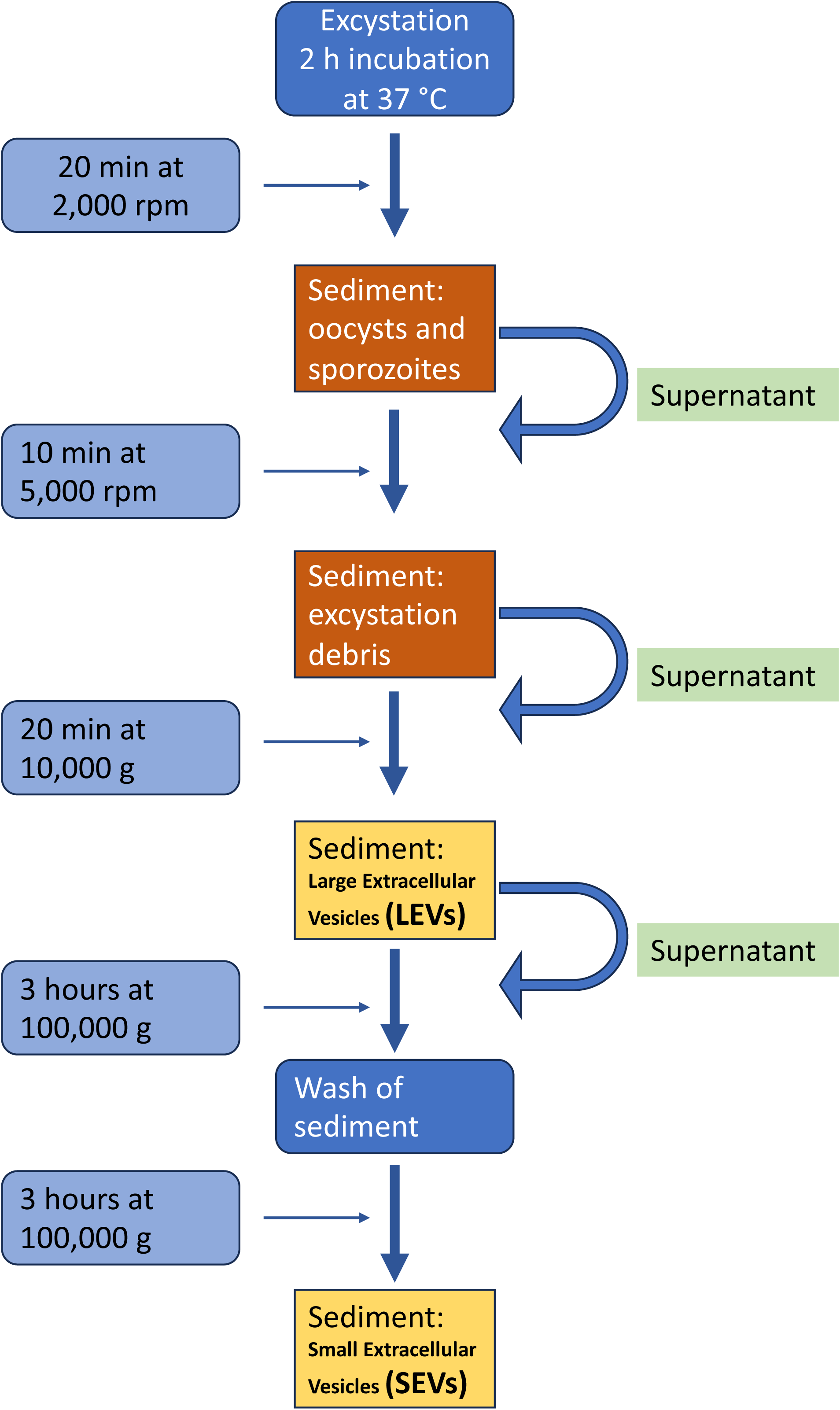
Diagram of the procedure used for the isolation of extracellular vesicles from the excystation medium.

### 2.4 Western blot analysis

Proteins in the excystation medium and supernatants after ultracentrifugation were precipitated with 10% ice-cold trichloroacetic acid (TCA)/0.015% (w/v) sodium deoxycholate (DOC). After 10 min on ice, samples were microfuged and the pellets were washed twice with 0.5 ml of ice-cold acetone, air dried, resuspended in 50 μl of lysis buffer (1% w/v SDS, 1% v/v Triton X-100, 0.5% w/v DOC, 10 mM 1,4-dithiothreitol (DTT), 1 mM EDTA) containing 1 μl/ml of Protease Inhibitor Cocktail (Sigma-Aldrich), and incubated for 5 min at 70 °C in ThermoMixer® (Eppendorf).

Pellets after centrifugation (LEVs) and ultracentrifugation (SEVs) were resuspended in 50 μl of lysis buffer containing 1 μl/ml of Protease Inhibitor Cocktail (Sigma-Aldrich) as above. Samples were boiled in Laemmli sample buffer before separation by SDS-PAGE on 4-20% TGX™ precast gels (Bio-Rad, Hercules, CA, USA). Gels were transferred to nitrocellulose membranes (Bio-Rad), which were then blocked in PBS with 5% non-fat milk in 0.1% Tween-20. Blots were incubated with primary antibodies, at 4 °C, overnight. For recombinant 6His-CpRom1 and 6His-CpGRASP, blots were incubated with mouse monoclonal RGS-His antibody (Qiagen) diluted 1:1,000. Blots for CpRom1 were probed with mouse anti-CpRom1 serum 1 diluted 1:250, while blots for CpGRASP were probed with mouse anti-CpGRASP serum diluted 1:100. For all blots incubation with secondary antibodies was conducted for 1 h, at 25 °C, with goat anti-mouse IgG-HRP conjugate (Bio-Rad, Hercules, CA USA) diluted 1:3,000. Detection of proteins was performed using Pierce ECL substrate (ThermoFisher) and bands were visualized with a ChemiDoc MP Imaging System (Bio-Rad, Hercules, CA, USA).

### 2.5 Ultrastructural analysis and immunolocalisation of extracellular vesicles by transmission electron microscopy

For negative staining in transmission electron microscopy (TEM), samples of LEVs and SEVs were suspended in 50 μl of 2.5% w/v glutaraldehyde in 0.1 M sodium cacodylate and incubated at 4 °C, overnight. Samples (20 μl) were deposited by successive applications on carbon-coated grids for electron microscopy; they were left to adsorb for 20 min, and the excess fluid was blotted with filter paper. A 2% w/v phosphotungstic acid contrasting solution (5 μl) was added on grids and air dried. Samples were observed by a PHILIPS Morgagni 268 TEM (FEI-Thermo Fisher) (Thery et al., 2006). For immunolocalization, LEVs and SEVs pellets were suspended in 100 μl of PBS, and aliquots of 20 μl were adsorbed on carbon-coated grids for electron microscopy, for 20 min. Samples were air dried and transferred to a drop of 2% w/v paraformaldehyde, for 5 min. After two washes with PBS, the grids were floated on PBS drops containing 0.1 M glycine for 30 min, washed with PBS, blocked with PBS containing 5% goat serum and 1% w/v BSA, for 30 min, and washed with PBS containing 0.05% w/v TWEEN 20 and 1% w/v BSA (PBS/BSA/TW). Then grids were floated on rabbit polyclonal anti-ID2 serum (1:10) in PBS/BSA/TW, for 1 h, at 25 °C, rinsed in the same buffer, incubated on 10 nm gold-conjugated goat anti-rabbit IgG (SIGMA) (1:50), for 1 h, rinsed again in buffer PBS/BSA/TW and treated with 1% glutaraldehyde, for 5 min. Finally, the samples were contrasted and embedded in a mixture 1:9 mixture of 4% uranyl acetate and 2% methyl cellulose respectively for 10 min on ice. The samples were air dried and analyzed by a PHILIPS Morgagni 268 TEM (FEI-Thermo Fisher) (Thery et al., 2006).

For ultrathin sections, 1 ml of excystation mixture (taken 30 min after induction) was loaded onto a 1 ml syringe and filtered with a 5 μm disposable filter Millex-SV (Millipore) to remove residual non-excysted and empty oocysts. Purified sporozoites were fixed in 2.5% w/v glutaraldehyde, 2% paraformaldehyde, 2 mM CaCl_2_ in 0.1 M sodium cacodylate, pH 7.4, and processed according to Perry and Gilbert (1979). Parasites were washed in cacodylate buffer and post-fixed with 1% OsO_4_ in 0.1 M sodium cacodylate, for 1 h, at 25 °C; then, they were treated with 1% tannic acid in 0.05 M cacodylate buffer, for 30 min, and rinsed in 1% sodium sulphate, 0.05 M cacodylate, for 10 min. Post-fixed specimens were washed, dehydrated through a graded series of ethanol solutions (30 to 100% ethanol) and embedded in Agar 100 (Agar Scientific Ltd, UK). Ultrathin sections, obtained by an UC6 ultramicrotome (Leica), were stained with uranyl acetate and Reynolds lead citrate and examined at 100 kV with a PHILIPS EM208S TEM (FEI - ThermoFisher) equipped with the Megaview III SIS camera (Olympus).

### 2.6 Scanning electron microscopy

1x10^7^ *C. parvum* oocysts (see above) were induced to excystation. At various times (from 0 to 30 min), the samples were fixed with 2.5% glutaraldehyde in 0.1 M sodium cacodylate buffer and 30 μl were left to adhere to polylysine-treated round glass coverslips (Ø 10mm), for 1 h, at 25 °C. Samples were processed for scanning electron microscopy (SEM) as previously described with slight modifications (Shively and Miller, 2009). Briefly, samples were post-fixed with 1% OsO_4_ in 0.1 M sodium cacodylate, for 1 h, at 25 °C, and were dehydrated through a graded series of ethanol solutions (from 30% to 100%). Then, absolute ethanol was gradually replaced by a 1:1 solution of hexamethyldisilazane (HMDS) and absolute ethanol, for 30 min, and successively by pure HMDS, for 1 h. Finally, HMDS was completely removed, and samples were left to dry in a desiccator, at room temperature, for 2 h, overnight. Then dried samples were mounted on stubs, carbon coated and analyzed by a FE-SEM Quanta Inspect F (FEI, Thermo Fisher Scientific).

### 2.7 Fluorescent labelling of extracellular vesicles and flow cytometric analysis

Fluorescent labelling of proteins of LEVs and SEVs was conducted as follows: pellets were resuspended in 50 μl PBS containing 10 µM Alexa Fluor 647 NHS Ester (Life Technologies) and incubated for 30 min, at 25 °C. Then pellets were resuspended again in 50 μl PBS containing 10 μM CFDA-SE (CFSE) (Life Technologies), for 30 min at room temperature. Reactions were stopped by adding 2 μl of 100 mM L-glutamine. Fluorescent LEVs and SEVs from the corresponding pellets were analysed by flow cytometry (FC) on a Gallios Flow Cytometer (Beckman Coulter) using an optimised procedure, as previously described (Coscia et al., 2016). Briefly, the instrument was set using the Flow Cytometry Sub-micron Particle Size Reference Kit providing green-fluorescent beads of different sizes (Invitrogen™) to establish the correct threshold value to apply on the 525/40 nm fluorescent channel (FL1). This procedure allows to create gates of the following dimensions: ≤200 nm, 500 nm and 1 µm. Sample analysis was performed by mixing 5 µl of fluorescent vesicles with 20 µl of Flow-Count Fluorospheres (Beckman Coulter) in a final volume of 200 µl of PBS to determine absolute counts. Fluorescent populations were analysed by plotting fluorescence at 525/40 nm (FL1) versus log scale side scatter (SSarea). Sample acquisition was stopped at 2,000 Flow-Count Fluorospheres events. The total number of fluorescent vesicles was established according to the formula: x = (((y × a)/b)/c) × d, where y = events counted at 2,000 counting beads; a = number of counting beads in the sample; b = number of counting beads registered (2,000); c = volume of sample analysed; and d = total volume of exosome preparation. Kaluza Software v. 2.0 (Beckman Coulter) was used for FC analysis.

### 2.8 Iodixanol gradient separation of extracellular vesicles

Freshly labelled fluorescent LEVs and SEVs from 1 x 10^7^ oocysts were diluted in 0.3 ml of PBS. Then, fluorescent samples were mixed with 1 ml of 60% iodixanol solution (OptiprepTM, Sigma-Aldrich), overlaid with 0.5 mL of 40%, 0.5 mL of 30% and 1.8 mL 10% of iodixanol solutions and floated into the gradient by ultracentrifugation in a SW60Ti rotor (Beckman) at 192,000 x g, for 18 hours, stopping without brake. After centrifugation, 12 fractions of 330 µl were collected from the top of the tube, diluted 40- to 80-fold with PBS and analysed by FC. Fraction densities were determined by refractometry. Gradient solutions were produced from the working solution (WS) by dilution with the HM solution. WS is prepared by mixing 5 volumes of OptiPrep™ with 1 volume of 0.25 M sucrose, 6mM EDTA, 60 mM Tris-HCl, pH 7.4; while HM solution contained 0.25 M sucrose, 1 mM EDTA, 10 mM Tris-HCl, pH 7.4.

Gradient fractions of fluorescent LEVs and SEVs were analysed by FC on a Gallios Flow Cytometer (Beckman Coulter) using an optimised procedure, as previously described (Coscia et al., 2016).

### 2.9 Proteomic analysis

Proteins from LEVs and SEVs were separated on a precast 4-15 % SDS-PAGE (Bio-Rad, Hercules, CA USA) and stained with Coomassie G-250. Gel lanes were cut into 9 slices and each one was *in-gel* reduced, alkylated with iodoacetamide and digested with trypsin, as previously reported (Salzano et al., 2013). Peptide mixtures were then desalted by μZipTipC18 tips (Millipore) before nano-liquid chromatography-electrospray ionization-Orbitrap tandem mass spectrometry (nLC-ESI-Q-Orbitrap-MS/MS) analysis, which was performed on a Q-Exactive Plus mass spectrometer equipped with UltiMate 3000 HPLC RSLC nano system (Thermo Fischer Scientific, USA). Peptides were separated on an Easy C18 column (100 × 0.075 mm, 3 μm) at a flow rate of 300 nl/min using a linear gradient from 5 to 40% acetonitrile in 0.1% formic acid, over 60 min. Mass spectra were acquired at nominal resolution 70,000 in the range m/z 350-1500, and data-dependent automatic MS/MS acquisition was applied to the ten most abundant ions (Top10), enabling dynamic exclusion with repeat count 1 and exclusion duration 30 s. The mass isolation window and the collision energy for peptide fragmentations were set to *m/z* 1.2 and 32%, respectively. Raw data from nLC-ESI-Q-Orbitrap-MS/MS analysis were searched by MASCOT v2.6.1, (Matrix Science, UK) node within Proteome Discoverer suite, against a *C. parvum* (strain Iowa II) database of protein sequences (4076 sequences) retrieved from CryptoDB site (https://cryptodb.org), and results were merged into a single mgf file to obtain proteins identified in LEVs and SEVs fractions. The following parameters were used for protein identification: mass tolerance values of 20 ppm and 0.05 Da for precursor and fragment ions, respectively; trypsin as proteolytic enzyme with maximum missed-cleavage sites of 2; Cys carbamidomethylation as fixed modification, Met oxidation, Asn/Gln deamidation and pyroglutamate formation from Gln/Glu, as variable modifications. A significance threshold of p<0.05 was set for protein identification, and molecular candidates with at least 2 significantly matched peptide sequences and protein Mascot score >50 were further considered for definitive assignment, after manual spectra visualization and verification.

### 2.10 Bioinformatic analyis

The sequence of CpRom1 (CryptoDB: cgd6_760) and CpGRASP (CryptoDB: cgd7_340) were downloaded at CryptoDB site (https://cryptodb.org). Similarity searches with DNA and protein sequences were conducted on non-redundant GenBank databases using the BLAST program (https://blast.ncbi.nlm.nih.gov/Blast.cgi). The prediction of transmembrane domains was performed at TMHMM-2.0 (http://www.cbs.dtu.dk/services/TMHMM/) and at Phobius (https://phobius.sbc.su.se/index.html). Prediction of structural and functional domains was performed at Pfam (http://pfam.xfam.org) and Prosite (https://prosite.expasy.org). The bidimensional representation of CpRom1 in Figure 2 was realized with TMRPres2D software (http://bioinformatics.biol.uoa.gr/TMRPres2D/download.jsp). Searches for potential cleavage sites by proteases were performed at PeptideCutter (https://www.expasy.org/resources/peptidecutter) and at ProP 1.0 server (http://www.cbs.dtu.dk/services/ProP/) for Arg and Lys propeptide cleavage sites (furin-like). Searches for potential GPI-anchor were performed at GPI Modification Site Prediction (https://mendel.imp.ac.at/gpi/gpi_server.html). Subcellular localization was predicted by DeepLoc-1.0 (http://www.cbs.dtu.dk/services/DeepLoc/) and was reported only when the probability was higher than 50%. Protein-protein interaction networks were obtained with STRING v.12.0 (https://string-db.org). Searches for proteins associated with extracellular vesicles were performed in the following databases: ExoCarta (http://www.exocarta.org) and Vesiclepedia (http://www.microvesicles.org).

**Figure 2.**
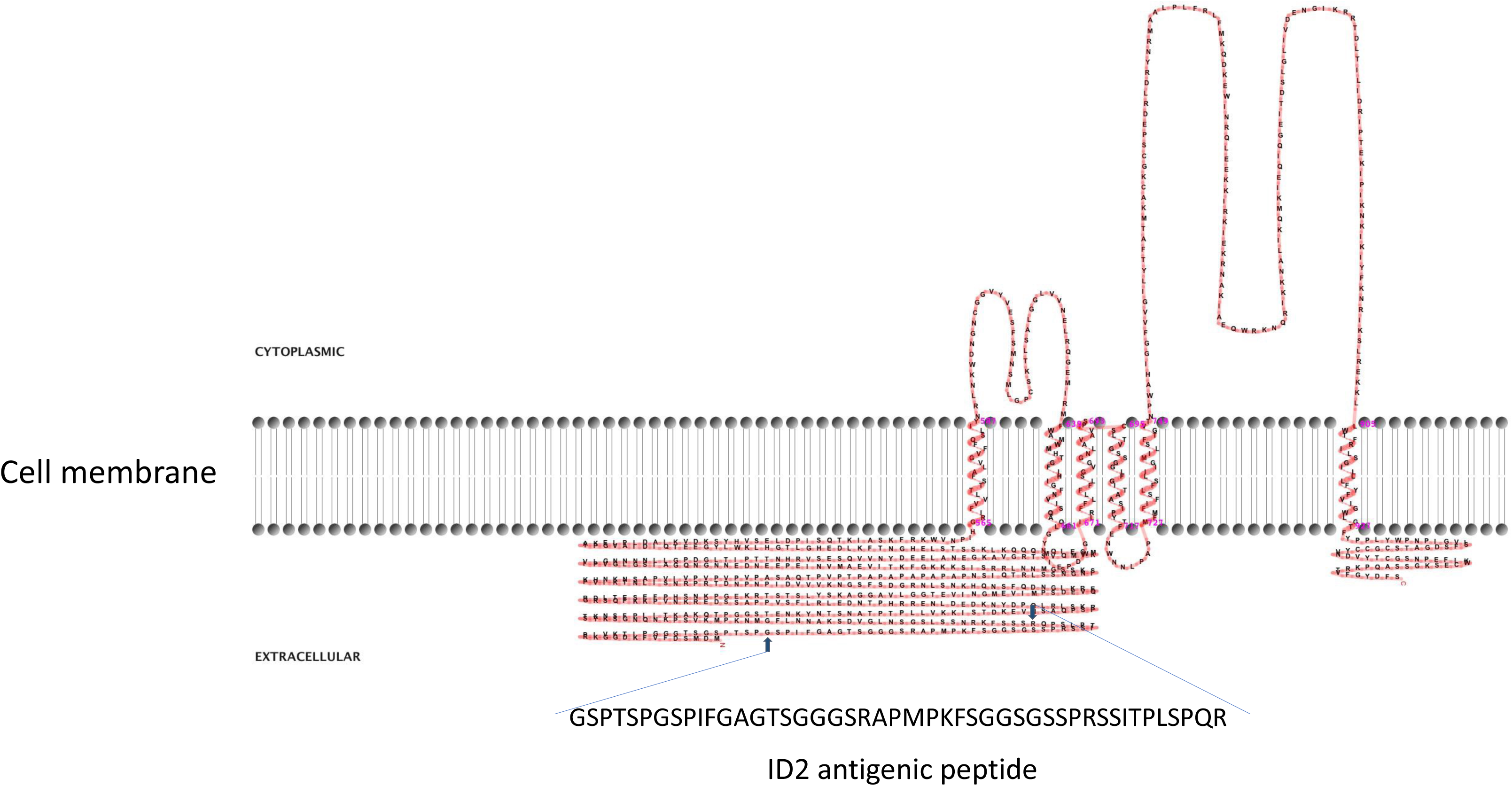
The two-dimensional model of CpRom1 in relation to the cell membrane: the diagram shows protein six transmembrane regions, two cytoplasmic loops (above the membrane), the large N-terminal extracellular region including the ID2 antigen, and the small extracellular region at the C-terminus (above the membrane). The small arrows show the position of the ID2 peptide (see text), and the amino acid sequence of the ID2 is expanded below.

## 3 Results

### 3.1 Characterization of CpRom1 expressed in oocysts and sporozoites

The CpRom1 rhomboid has been previously identified starting from the 46 amino acid-long ID2 peptide (Trasarti et al., 2007). A prediction based on the protein sequence showed that CpRom1 consists of 990 residues with a theoretical molecular mass of 109.3 kDa; this makes CpRom1 the largest rhomboid ever described. Computational predictions showed that CpRom1 has a large N-terminal portion of approximately 500 amino acids, which includes the ID2 peptide (Figure 2). This large portion of the protein is exposed towards the extracellular environment and does not show any recognizable domain (figure 2). A well-distinguishable rhomboid domain, which consists of six transmembrane helices surrounding the proteolytic site, is contained in the final 450 amino acids at the C-terminus.

To identify the native CpRom1 rhomboid, we prepared a mouse polyclonal serum reacting with the full-length protein. To this end, a fusion protein with six histidine-tag (6His-CpRom1) was expressed in bacteria, purified by Ni-NTA chromatography and used to immunize Balb-C mice. The polyclonal mouse antiserum was used in Western blot experiments on sporozoite lysates, in which a band of approximately 110 kDa was recognized, in agreement with an expected molecular mass of 109.3 kDa (Figure 3). However, the serum also detected a smaller band migrating at about 75 kDa, which might represent a cleaved form of the 110 kDa protein. These CpRom1 forms were present in resting oocyst at time 0 of excystation (Figure. 3A), with a similar amount of the 110 kDa form in unexcysted oocyst and sporozoites, whereas a lower amount of the 75 kDa species was detected after excystation. Probably, the latter protein form is quickly degraded after excystation. A significant amount of CpRom1 was also detectable in the supernatant of excystation medium after eliminating sporozoites and residual oocysts. In fact, TCA precipitates of excystation medium after 30 min of incubation, probed with anti-CpRom1 serum revealed a distinctive band of 110 kDa (Figure 3B). Given the presence of six transmembrane domain in CpRom,1 which eventually hampers its release in the excystation medium, we hypothesized that CpRom1 is released in association with membranous vesicles.

**Figure 3.**
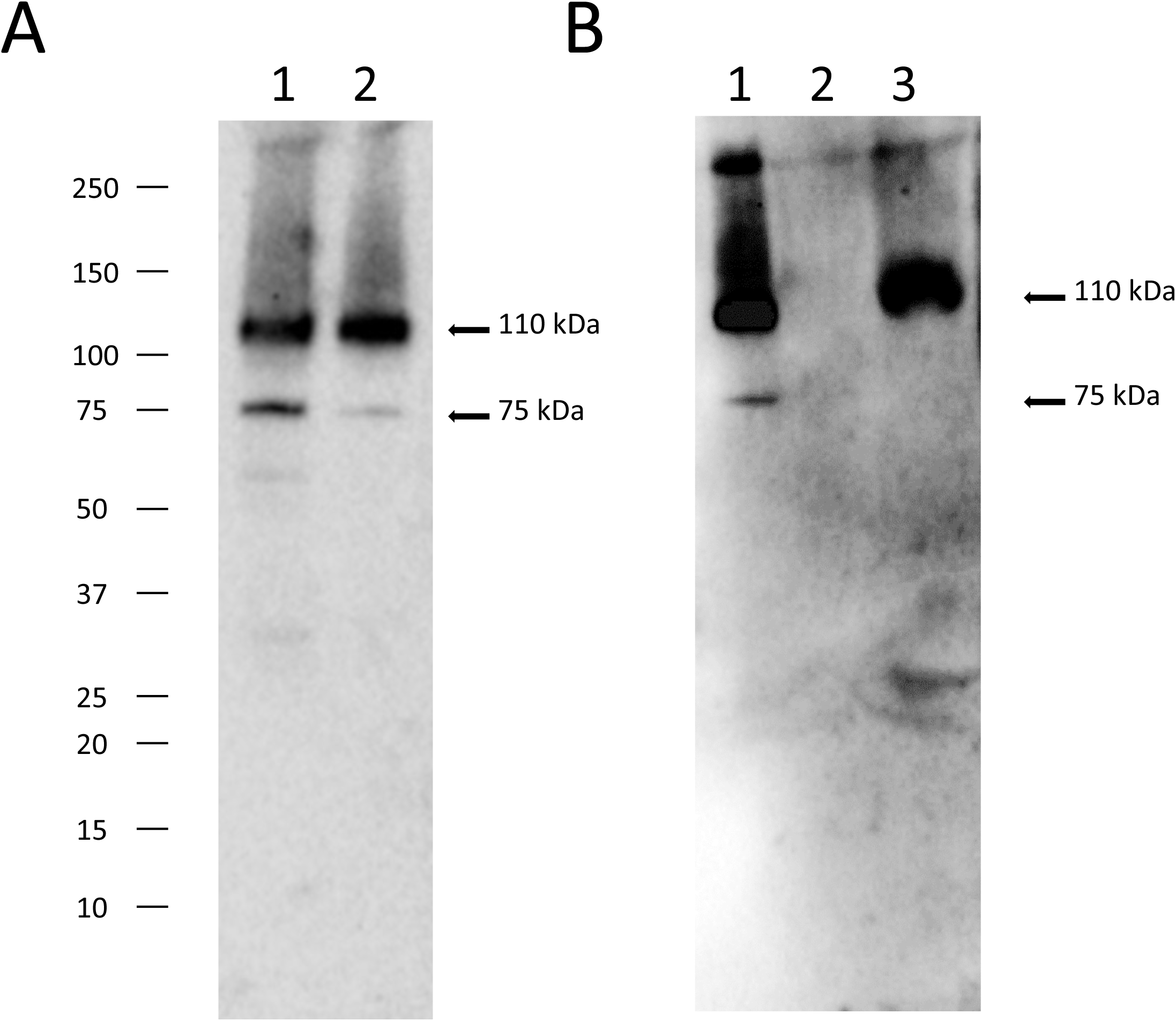
CpRom1 identification by Western blot. A, Western blotting on oocysts and sporozoite lysates probed with mouse anti-CpRom1 serum: 1, lysate from 5x10^6^ unexcysted oocysts; 2, lysate from 5x10^6^ oocysts after 30 min of excystation. B, Western blotting on excystation medium and TCA-precipitated supernatant of probed with mouse anti-CpRom1 serum: 1, lysate from 1x10^7^ oocysts after 30 min of excystation; 2, TCA-precipitated supernatant before the excystation; 3, TCA-precipitated supernatant of sample 1.

### 3.2 CpRom1 is associated with released extracellular vesicles during the excystation

To test this the hypothesis reported above, we developed an ultracentrifugation protocol to separate extracellular vesicle populations from the excystation medium, after preventive removal of oocyst-sporozoite residues. A schematic representation of the procedure is shown in Figure 1. Depending on centrifugation conditions, two types of *C. parvum* EVs were selectively recovered that were tentatively classified as large extracellular vesicles (LEVs) and small extracellular vesicles (SEVs). Protein extracts from LEVs and SEVs were then assayed for the presence of CpRom1 by dedicated immunoblotting. Results showed that a protein band of 110 kDa was observed in the LEVs extracts, but not in the SEVs counterparts (Figure 4A).

**Figure 4.**
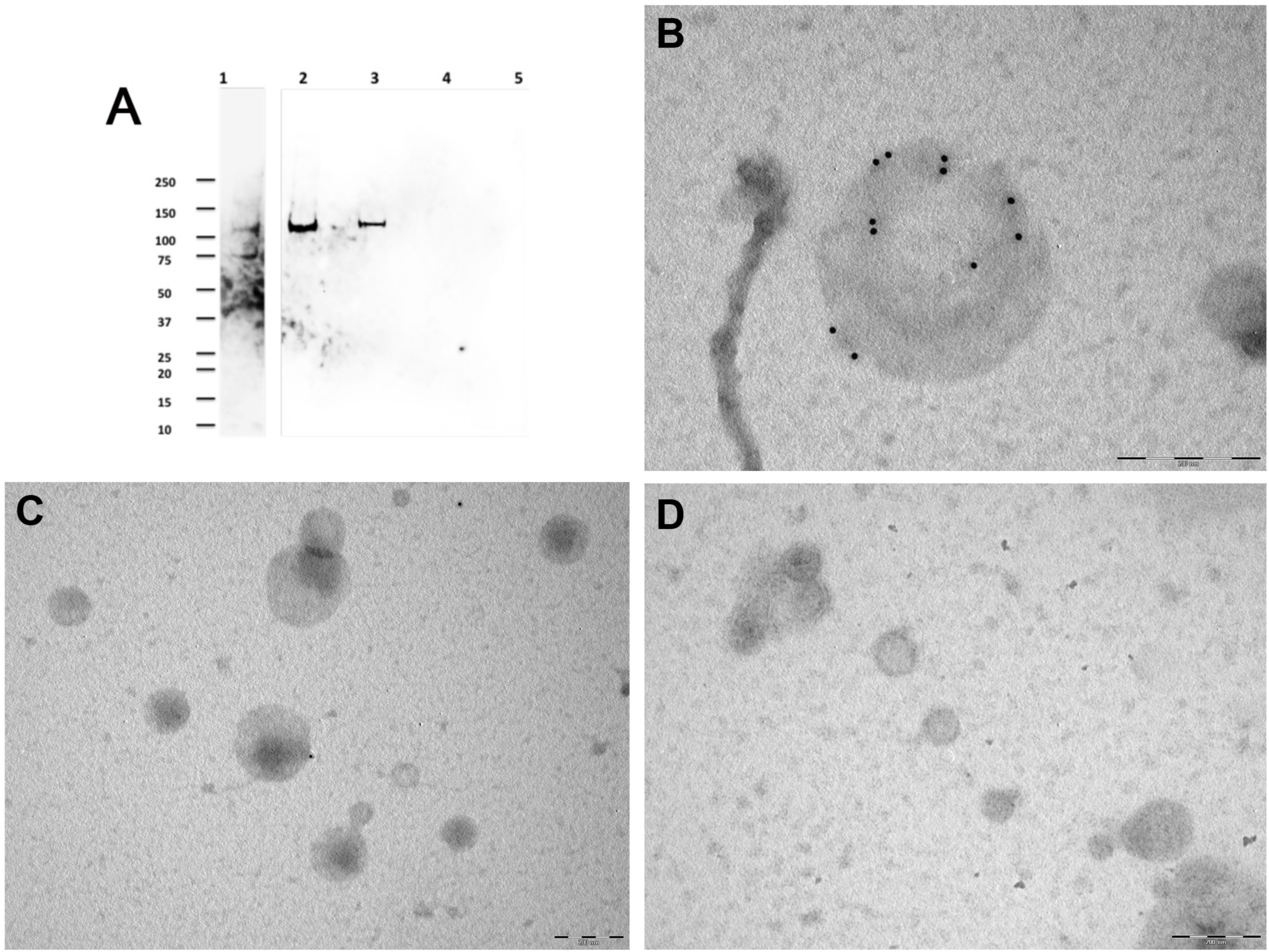
Immunolocalization of CpRom1 by Western blot on microvesicle extracts and immunoelectron microscopy of the corresponding extracellular vesicles. A, Immunoblotting with anti-CpRom1 mouse serum: 1, unexcysted oocysts lysate; 2, excysted soporozoites; 3, LEVs extract; 4, SEVs extract; 5, TCA-precipitated supernatant of SEVs. B, Negative staining immunoelectron microscopy of LEVs labelled with anti-CpRom1 rabbit serum. C, Negative staining immunoelectron microscopy of LEVs labelled with pre-immune rabbit serum. D, Negative staining immunoelectron microscopy of SEVs labelled with anti-CpRom1 rabbit serum.

To demonstrate that CpRom1 is associated with EVs, we assayed *C. parvum* LEVs and SEVs by negative staining immunoelectron microscopy using a polyclonal rabbit serum directed against the ID2 peptide (Trasarti et al., 2007). In agreement with the immunoblotting results, numerous strong immunogold signals were detected on the surface of large vesicles (> 150 nm) occurring in LEVs (Figure 4B). A negative control with pre-immune rabbit serum showed sporadic signals with one or two gold particles for microscopic field (Figure 4C). We also tested SEVs by negative staining immunoelectron microscopy with the same anti-ID2 serum, and we observed small vesicles that were not labelled by immunogold particles (Figure 4D).

All together, these results suggested that sporozoites release two different types of extracellular vesicles, among which those present in LEVs are generally larger than those in SEVs. Immunolabelling for CpRom1 showed the presence of this protein only in LEVs.

### 3.3 Characterization of sporozoite extracellular vesicles by flow cytometry and electron microscopy

To further characterize sporozoite EVs, we labelled vesicles in LEVs and SEVs sequentially with two different fluorescent probes: at first, we used the hydrophilic dye NHS-AF647, which labels proteins on the external surface of the vesicles; then, we utilized with the lipophilic dye CFSE, which passively diffuse into vesicles labelling also internal vesicular proteins. Successively, vesicles contained in LEVs and SEVs were separated by density gradient centrifugation on an iodixanol gradient, and the fluorescent content of fractions was analysed by flow cytometry. FC analysis showed that LEVs comprised a heterogeneous population of intact vesicles ranging in size from 50 nm to over 250 nm (Figure 5A, B). Notably, LEVs were often clumped together (Figure 6A), contributing to the highest density of extracellular vesicles according to FACS analysis. In contrast, SEVs displayed a more homogeneous composition (Figure 5C), consisting of smaller intact vesicles with sizes below 150 nm (Figure 6B).

**Figure 5.**
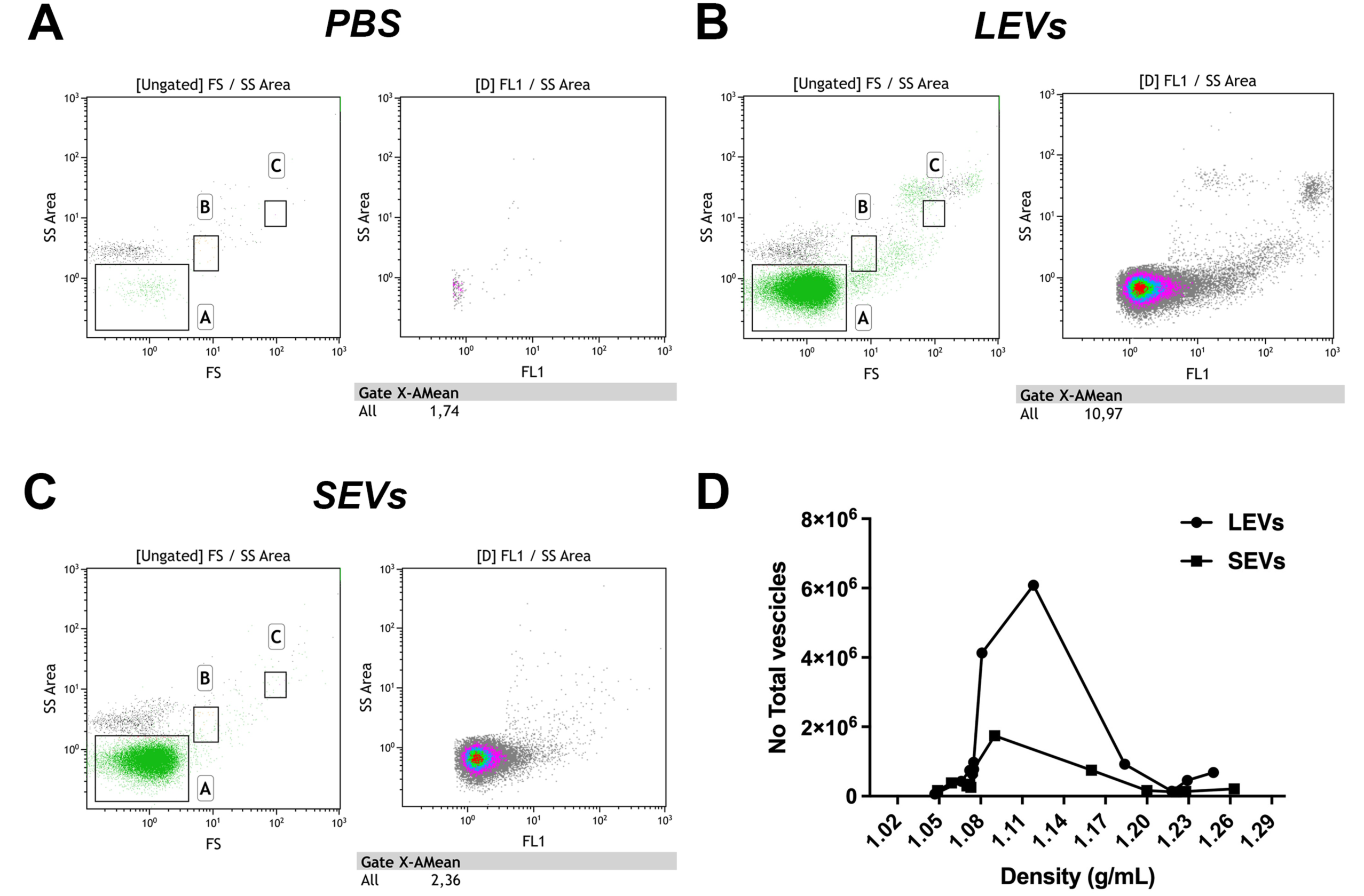
Flow cytometry analysis of gradient fractions of LEVs and SEVs. A, PBS buffer used to resuspend EVs as negative control. B, Dot plot of LEVs fraction at 1.1 g/ml density. C, Dot plot of SEVs at 1.1 g/ml density. D, diagram comparing the distribution of LEVs and SEVs in the gradient fractions. Left panels indicate gate dimensions (A≤200 nm, B=500 nm and C=1 µm). Right dot plot panels show vesicles distribution in terms of fluorescence.

**Figure 6.**
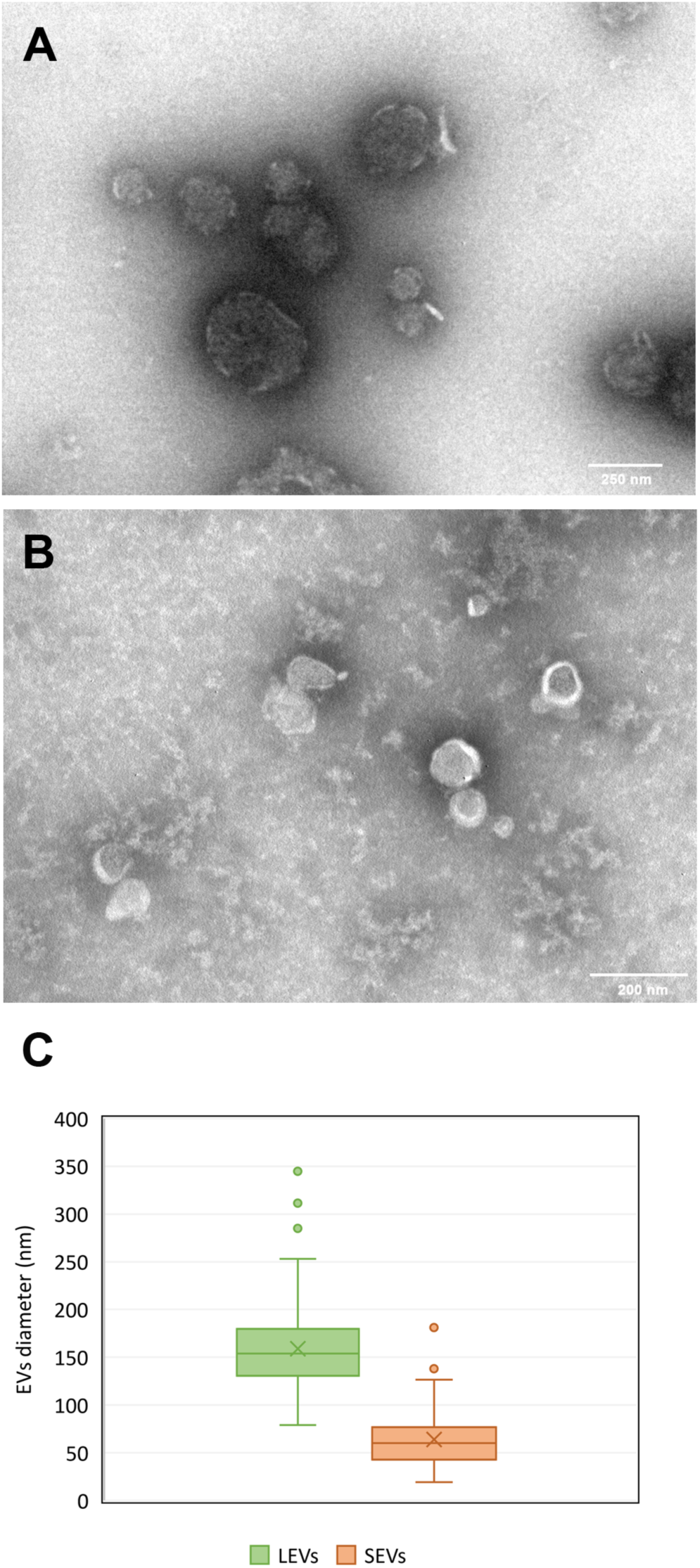
Physical characterization of LEVs and SEVs by electron-microscopy. A, TEM negative staining of LEVs. B, TEM negative staining of SEVs. C, diagram showing percentage distribution of vesicle size in LEVs (green) and in SEVs (orange).

EVs were also quantified by FC. We observed that LEVs are more numerous than SEVs; moreover, both vesicle populations presented a peak in fractions with density ranging between 1.08-1.14 g/ml (Figure 6D). To establish more specifically the size of vesicles occurring in LEVs and SEVs, the diameters of a hundred vesicles were measured through electron microscopy, and the corresponding micrographs were subjected to statistical analysis. This experiment revealed that LEV (figure 6A) and SEV (figure 6B) had a different medium size of 150 nm and 60 nm respectively. (Figure 6C).

### 3.4 Ultrastructural characterization of vesicles during the excystation

To capture images of vesicles at the time of their release from sporozoites, we analysed by SEM and TEM the excystation mixture at various time points (from 0 to 30 min) after the induction. Hence, we observed some sporozoites immediately after the egress from two oocysts (Figure 7A). Oocysts appeared as thin, empty envelopes, and oocysts and sporozoites are surrounded by numerous vesicles of varying sizes scattered throughout the microscopic field. Similarly, we captured images at higher magnification showing two vesicles of different sizes at the very moment of their release from the sporozoite membrane (Fig. 7B). Finally, we observed emerging vesicles from the apical part of a sporozoite (Figure 7C) as well as a vesicle of approximately 200 nm just released from the posterior part of another sporozoite (Figure 7D). Together, these images supported the origin of EVs, at least the larger ones, by budding of the external membrane.

**Figure 7.**
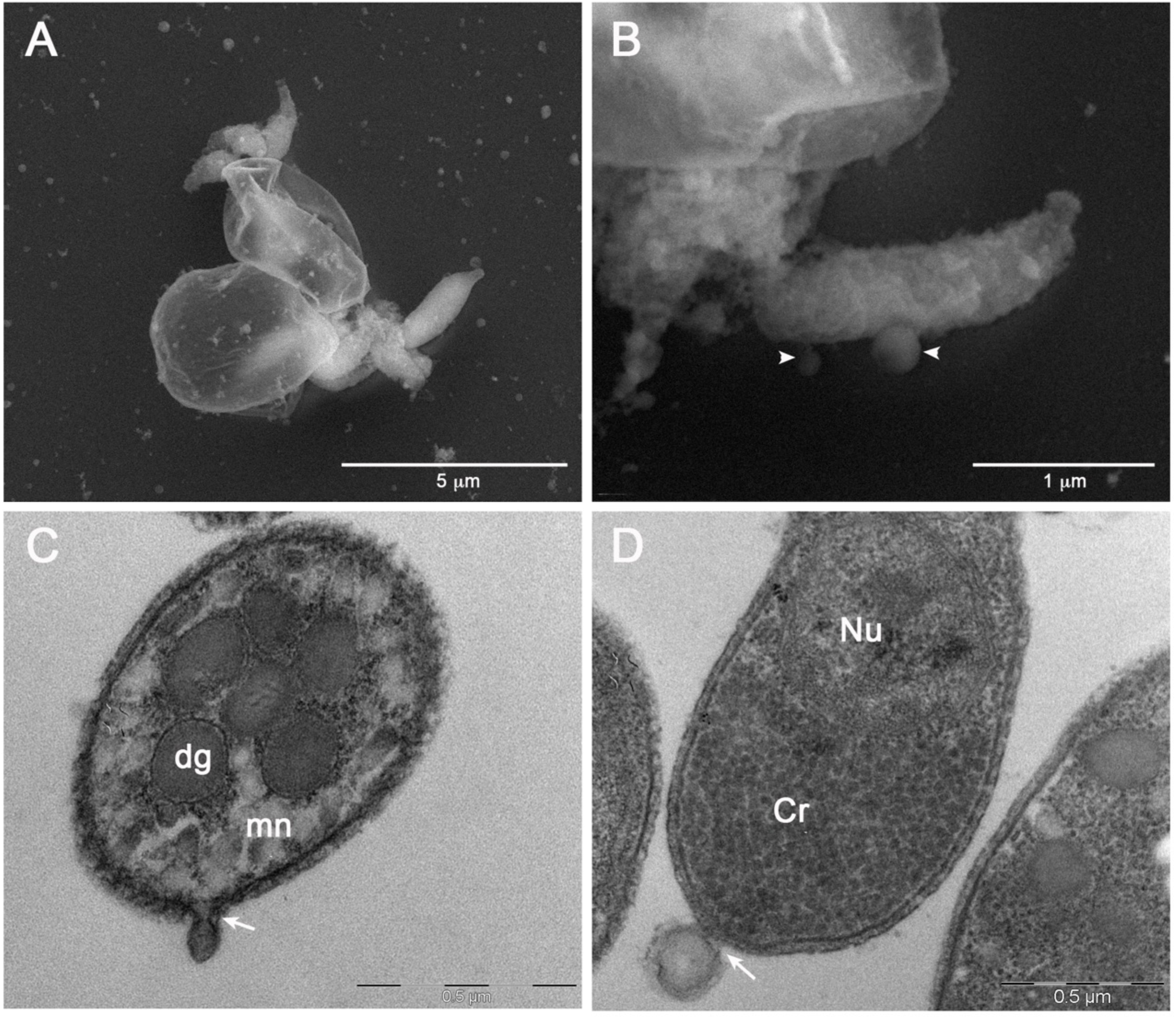
Electron microscopy images of EVs at their release from sporozoites. A, Backscattered electron SEM micrograph of two excysted oocysts showing the release of vesicles during the sporozoites egress. B, High magnification of an egressed sporozoite which shows two budding vesicles (head arrows). C, TEM micrograph showing the plasmamembrane budding of a vesicle from the apical region of a sporozoite (white arrow). D, TEM micrograph of a vesicle budding from the posterior region of a sporozoite. DG: dense granules; mn: micronemes; Cr: crystalloid; Nu: nucleus.

Since SEVs have physical parameters like exosomes, which originate from intracellular multi-vesicular bodies (MVBs) (Van Niel et al., 2018; Barreca et al., 2023), we looked for a similar structure in the sporozoite cytoplasm. Electron microscopy images showed a vacuolar structure with internal membranes that resemble MVBs (Figure 8A and 8B). A similar MVB-like structure has not yet been described so far in *Cryptosporidium* spp.

**Figure 8.**
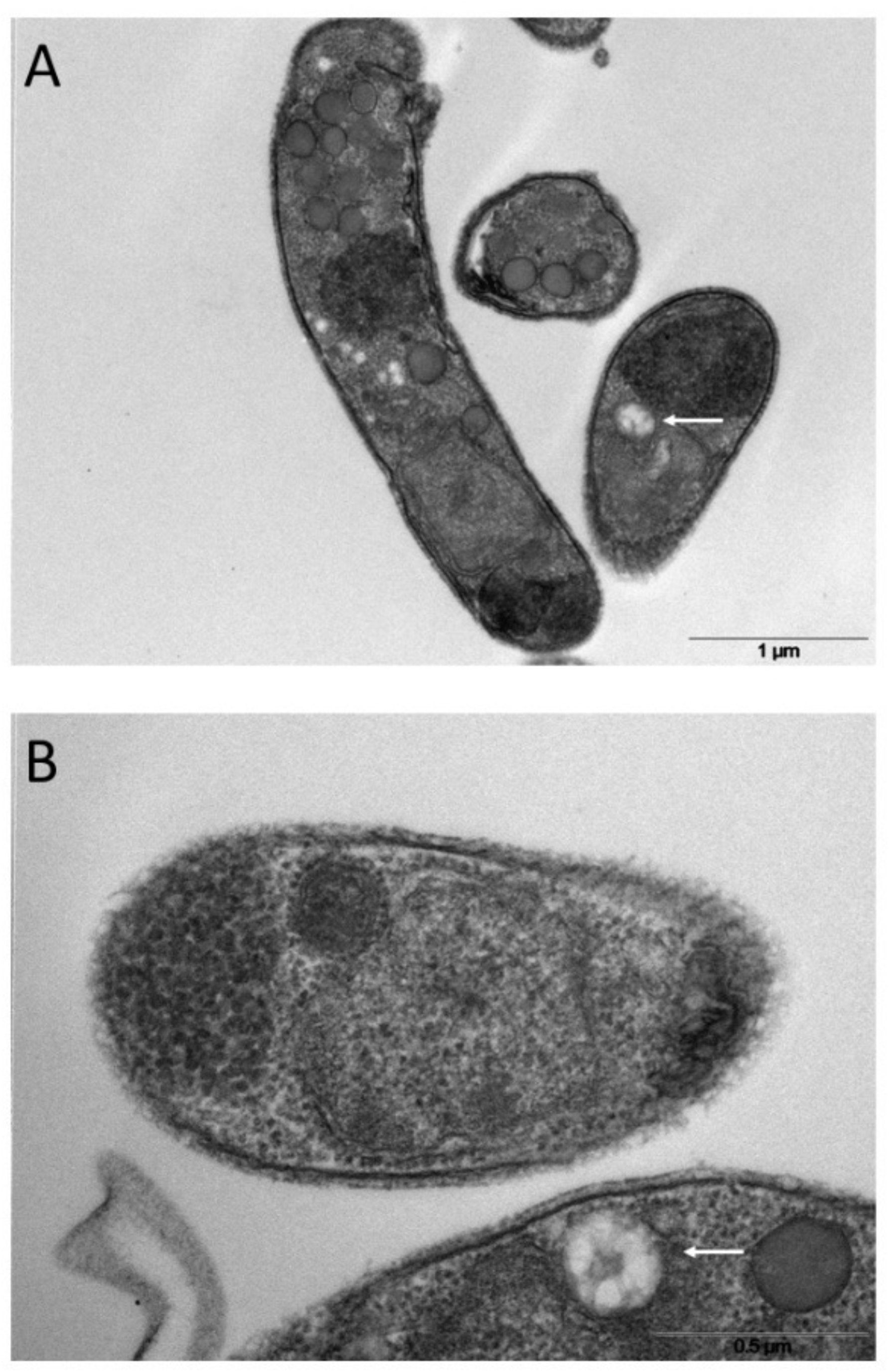
Ultrathin sections of excysted sporozoites (A, B) showing possible MVB-like organelles inside the cells (white arrows).

### 3.5 The Golgi protein CpGRASP is associated with LEVs but not with SEVs

No homologues of the most common mammalian proteins associated with exosomes, namely tetraspanins (eg, CD63, CD81 or CD9) or endosomal sorting complex required for transport (ESCRT) proteins (eg, Alix and Tsg101) (Théry et al. 2018), can be identified in *Cryptosporidium* proteome (data not shown). On the other hand, well-conserved *Cryptosporidium* homologues of proteins involved in vesicle trafficking can be traced back among those associated with the Golgi apparatus. In all eukaryotes, the Golgi complex is a central hub for the vesicle trafficking (Doyle and Wang, 2019), and Golgi proteins could represent markers for tracking various types of vesicles. The Golgi reassembly and stacking proteins (GRASPs) belong to a protein family with a conserved similarity in all eukaryotes, and the *Cryptosporidium* homolog is easily identifiable in CryptoDB. Moreover, following a search in protein databases of extracellular vesicles and in the literature, GRASPs have been found in different types of EVs (Chua et al., 2012; Peres da Silva et al., 2018). Therefore, we proceeded to clone the gene for the *C. parvum* homolog of GRASPs to express a histidine-fusion protein in bacteria. The purified recombinant protein (6his-CpGRASP) was then used to generate a specific antiserum in mice. In Western blot experiments, when probed with the anti-GRASP serum, an 80 kDa band was identified (Figure 9) in the whole oocyst-sporozoite lysate, which corresponded to the expected molecular mass of 82.49 kDa. The same 80 kDa band was also observed in LEVs extract, while no band was detected in the SEVs extract or in the supernatant following the ultracentrifugation. This last result suggested that the biogenesis of LEVs involves Golgi-derived vesicles unlike SEVs, which conversely did not show any evidence for thre occurrence of CpGRASP.

**Figure 9.**
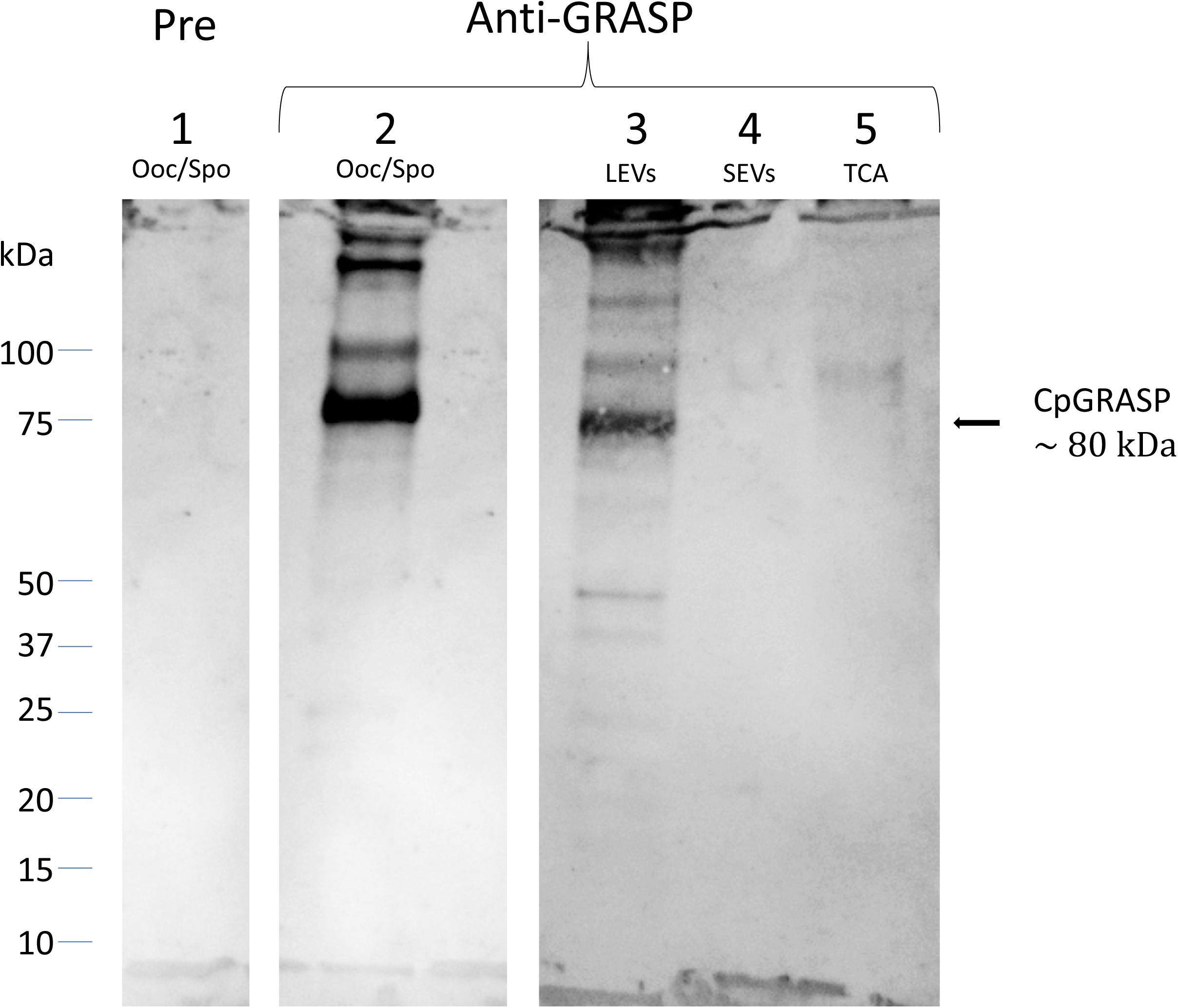
Western blot probed with mouse anti-CpGRASP serum. 1, lysate oocyst-sporozoite probed with pre-immune serum; 2, lysate oocyst-sporozoite; 3, LEVs extract; 4, SEVs extract; 5, SEVs supernatant precipitated with TCA. 4-20% SDS-PAGE, lane 1 probed with 1:500 with mouse serum before the immunization; lane 2-5 probed with 1:500 mouse serum after the immunization with recombinant CpGRASP.

### 3.6 Comparison of protein profiles of LEVs and SEVs by fluorescent labelling and SDS-PAGE electrophoresis

A first analysis regarding the protein content in sporozoite vesicles was performed by comparing the electrophoretic profiles of the two types of vesicles. To this end, we double-labeled the LEVs and SEVs with NHS-AF647 (red) and CFSE (green) and analysed the corresponding protein profiles by SDS-PAGE (Dehghani and Gaborski, 2020). Figure 10 shows the image obtained by overlaying the green and red channels onto the electrophoretic protein profiles of LEVs and SEVs. Upon comparing the electrophoretic profiles, we have identified distinct common bands (black arrows) as well as some prominent unique bands (white arrows) in SEVs. This method allowed a rapid generation of distinctive patterns for LEVs and SEVs and confirmed the occurrence of differences in the protein content of them. Of note, the SEVs profile exhibited a greater number of red bands indicating the likely presence of a greater number of surface proteins.

**Figure 10.**
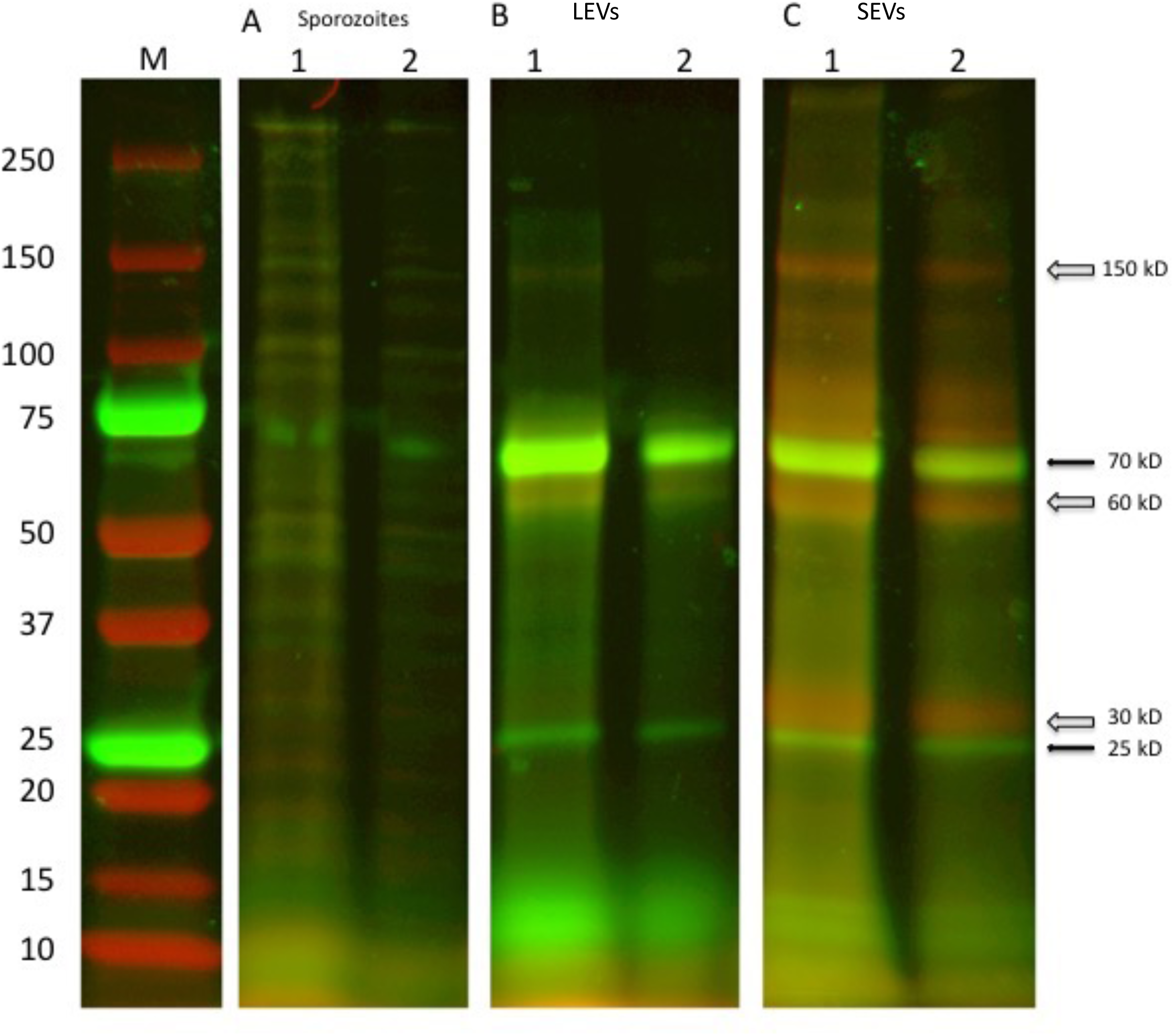
SDS-PAGE of two independent excystation experiments followed by labelling with the fluorescent dyes NHS-AF647 (red) and CFSE (green) of the different stages of centrifugation: 1 with 1x10^7^oocysts, 2 with 5 x 10^6^ oocysts duplicate sporozoite lysate; LEVs and SEVs after labelling with NHS-AF647 (red) and CFSE (green) molecular probes. Black arrows indicate common green bands, open arrows indicate red bands present only in SEVs extract.

### 3.7 Proteomic analysis of LEVs and SEVs content

To identify *C. parvum* proteins present in above-reported vesicles, components of LEVs and SEVs were resolved by SDS-PAGE and stained with Coomassie G250. The resulting whole gel lanes (see Supplemental Figure S1) were divided into 9 slices that were further subjected to trypsinolysis and then to nLC-ESI-Q-Orbitrap-MS/MS analysis. On the whole, we identified 60 *C. parvum* proteins of which 39 were in common between the two vesicle types; conversely, 5 proteins were identified only in LEVs, while 16 were found exclusively in SEVs. All identified proteins are listed in Table 1, and details of the corresponding proteomic analysis are reported in Supplemental Table S1. The full annotation from the CriptoDB database for all assigned proteins is reported in Supplemental Table S2, which also includes presumptive information on the corresponding function, gene ontology (GO terms ID), assignment as membrane/soluble component, and cellular localisation.

**Table 1.** List of identified proteins in C. parvum EVs by proteomic experiments. Proteins in the light blue background were found both in LEVs and SEVs, proteins in yellow background were found only in LEVs, and proteins in green background only in SEVs.

| Protein ID (CryptoDB) | Gene product (CryptoDB) | MW (kDa) | GO terms (CryptoDB) -Presumptive function and/or localization | <u>Evs DB[1]</u> | LEVs | SEVs |
| --- | --- | --- | --- | --- | --- | --- |
| cgd1 3020-RA | Fructose-bisphosphate aldolase | 38.5 | glycolytic process | √ | y | y |
| cgd1 3040-RA | Triosephosphate isomerase | 27.4 | glycolytic process gluconeogenesis | √ | y | y |
| cgd1 870-RA | Peptidyl-prolyl cis-trans isomerase | 22.9 | protein peptidyl-prolyl isomerization/protein folding | √ | y | y |
| cgd2 20-RA | Heat shock-70 protein | 73.3 | No Data | √ | y | y |
| cgd2 3330-RA | Hsp70-protein | 103 | ATP binding | √ | y | y |
| cgd2 3950-RA | Translation-elongation factor EF1B/ribosomal protein S6 | 27.6 | translational elongation | √ | y | y |
| cgd2 4320-RA | Thioredoxin / glutathione reductase selenoprotein | 56.3 | cell redox homeostasis | √ | y | y |
| cgd3 1290-RA | 14-3 -3 domain containing protein | 28.6 | No Data | √ | y | y |
| cgd3 1540-RA | Signal peptide containing protein | 185 | No Data | - | y | y |
| cgd3 2090-RA | 40S ribosomal protein Sae | 28.3 | Translation /Ribosome | √ | y | y |
| cgd3 3370-RA | Uncharacterized protein | 166 | No Data | - | y | y |
| cgd3 3430-RA | Amine oxidase | 199 | amine metabolic process /copper ion binding | √ | y | y |
| cgd3 3770-RA | Hsp90 protein | 80.8 | protein folding / ATP binding | √ | y | y |
| cgd4 1940-RA | Nucleoside diphosphate kinase | 16.8 | nucleoside diphosphate phosphorylation | √ | y | y |
| cgd4 2600-RA | UDP-glucose 4-epimerase | 38.5 | galactose metabolic process / UDP-glucose 4-epimerase activity | √ | y | y |
| cgd4 3270-RA | Hsp70 protein | 90.7 | ATP binding | √ | y | y |
| cgd4 740-RA | Thioredoxin peroxidase-like protein | 21.8 | peroxidase activity | √ | y | y |
| cgd5 1960-RA | Enolase | 48.4 | glycolytic process / magnesium ion binding | √ | y | y |
| cgd5 2020-RA | Cysteine-rich secretory protein. Allergen V5/Tpx-1-related | 46.2 | No Data | - | y | y |
| cgd5 2800-RA | Actin depolymerizing factor | 15.5 | actin filament depolymerization / actin cytoskeleton / actin binding | √ | y | y |
| cgd5 3160-RA | Actin | 42.1 | No Data | √ | y | y |
| cgd5 3360-RA | Adenylate kinase | 24.2 | adenylate kinase activity | √ | y | y |
| cgd5 3740-RA | 50S ribosomal protein L7e/L30e/S12e/Gadd45 | 15.8 | translation/Ribosome | √ | y | y |
| cgd6 1080-RA | Glycoprotein GP40 | 33.4 | No Data | - | y | y |
| cgd6 120-RA | Disulfide-isomerase. Signal peptide plus ER retention motif | 53.8 | endoplasmic reticulum lumen / protein disulfide isomerase activity | √ | y | y |
| cgd6 2450-RA | Glycogen / starch / alpha-glucan phosphorylase | 104 | carbohydrate metabolic process / glycogen phosphorylase activity | √ | y | y |
| cgd6 2690-RA | FKBP-like peptidyl-prolyl isomerase | 36.8 | protein folding/peptidyl-prolyl cis-trans isomerase activity | √ | y | y |
| cgd6 3790-RA | Glyceraldehyde-3-phosphate dehydrogenase | 36.1 | glycolytic process / glyceraldehyde-3-phosphate dehydrogenase (NAD+) (phosphorylating) activity | √ | y | y |
| cgd6 4460-RA | Uncharacterized protein with Armadillo-like helical | 276 | No Data | - | y | y |
| cgd6 710-RA | Uncharacterized Secreted Protein | 49.7 | No Data | - | y | y |
| cgd7 1710-RA | Threonyl-tRNA synthetase | 87.6 | tRNA aminoacylation/cytoplasm/aminoacyl-tRNA ligase activity | √ | y | y |
| cgd7 3670-RA | Heat shock protein Hsp90 | 89.1 | protein folding / ATP binding | √ | y | y |
| cgd7 400-RA | Uncharacterized protein | 38.6 | No Data | - | y | y |
| cgd7 4310-RA | Cysteine-rich secretory protein. Allergen V5/Tpx-1-related | 185 | No Data | - | y | y |
| cgd7 4450-RA | Elongation factor EF1-gamma (Glutathione S-transferase family) | 43.1 | translational elongation / transferase activity | √ | y | y |
| cgd7 480-RA | L-lactate / malate dehydrogenase | 33.9 | carbohydrate metabolic process / oxidoreductase activity | √ | y | y |
| cgd7 910-RA | Phosphoglycerate kinase | 42 | glycolytic process / phosphoglycerate kinase activity | √ | y | y |
| cgd8 1720-RA | Aldehyde / Alcohol dehydrogenase | 89.7 | alcohol dehydrogenase (NAD+) activity | √ | y | y |
| cgd8 2930-RA | Gtpase translation elongation factor 2 | 92.7 | GTPase activity | √ | y | y |
| cgd1 2040-RA | Pyruvate kinase | 56.4 | glycolytic process / pyruvate kinase activity | √ | y | n |
| cgd1 660-RA | Signal peptide region containing protein | 37.1 | No Data | - | y | n |
| cgd4 2300-RA | Ubiquitin-activating enzyme E1 | 120 | cellular protein modification process / ubiquitin-like modifier activating enzyme activity | √ | y | n |
| cgd7 2280-RA | Ribosomal protein L40e | 14.7 | translation / ribosome / structural constituent of ribosome | √ | y | n |
| cgd7 4270-RA | 2,3-bisphosphoglycerate-dependent phosphoglycerate mutase | 28.3 | glycolytic process/phosphoglycerate mutase activity | √ | y | n |
| cgd1 2860-RA | UbiE/COQ5 methyltransferase | 32.4 | methylation/methyltransferase activity | √ | n | y |
| cgd1 3690-RA | Aspartyl (Acid) protease | 88.4 | proteolysis/integral component of membrane/aspartic-type endopeptidase activity | √ | n | y |
| cgd2 3780-RA | Uncharacterized protein | 58.9 | No Data | √ | n | y |
| cgd2 4120-RA | Peptidyl-prolyl cis-trans isomerase | 18.5 | protein folding / peptidyl-prolyl cis-trans isomerase activity | √ | n | y |
| cgd2 860-RA | Proteasome subunit beta type | 22.9 | proteolysis involved in cellular protein catabolic process /proteasome core complex /threonine-type endopeptidase activity | √ | n | y |
| cgd5 1650-RA | DJ-1/Pfpl | 19.6 | No Data | √ | n | y |
| cgd5 3230-RA | Manganese/iron superoxide dismutase | 25.5 | superoxide metabolic process / superoxide dismutase activity | √ | n | y |
| cgd5 440-RA | Adenylate cyclase-associated CAP | 19 | cytoskeleton organization/actin binding | - | n | y |
| cgd6 1630-RA | CS domain containing protein | 21.2 | No Data | √ | n | y |
| cgd6 3180-RA | 40S ribosomal protein S15 | 16.4 | translation/ribosome/small ribosomal subunit/structural constituent of ribosome | - | n | y |
| cgd6 3970-RA | Glutaredoxin-like protein 2 thioredoxin folds | 24.7 | protein disulfide oxidoreductase activity / iron-sulfur cluster binding | √ | n | y |
| cgd6 4320-RA | 40S ribosomal protein S5 | 21.8 | translation / small ribosomal subunit / structural constituent of ribosome | √ | n | y |
| cgd7 220-RA | GTP-binding nuclear protein | 24.1 | nucleocytoplasmic transport / nucleus / GTPase activity | √ | n | y |
| cgd7 3120-RA | Thiamine pyrophosphate enzyme | 64.4 | carboxy-lyase activity / thiamine pyrophosphate binding | √ | n | y |
| cgd7 360-RA | Heat shock protein 70 | 71.8 | endoplasmic reticulum / ATP binding | √ | n | y |
| cgd8 2110-RA | MIR motif-containing 39-like glycosyltransferase | 25.4 | membrane | √ | n | y |

[1] Presence of related proteins in Vesiclepedia (http://microvesicles.org) or ExoCarta (http://exocarta.org): √

According to functional predictions, the largest number (14) of annotated proteins included enzymes related to cell metabolism, while the second group (12 in number) consisted of components involved in protein folding, such as heat shock proteins (HSPs) and molecular chaperones (Figure 11, panel A). This latter group included 4 heat shock proteins 70 (HSP70s) and 2 heat shock proteins 90 (HSP90s) that may have a different subcellular localization, as inferred by the variable presence of signals and/or transmembrane domains. Additional protein categories were related to protein synthesis (9 in number), regulatory function (6 in number), cytoskeleton (3 in number), redox homeostasis (2 in number) and proteolysis (2 in number) (Figure 10, panel A). The presumptive function remained unknown for 12 proteins.

**Figure 11.**
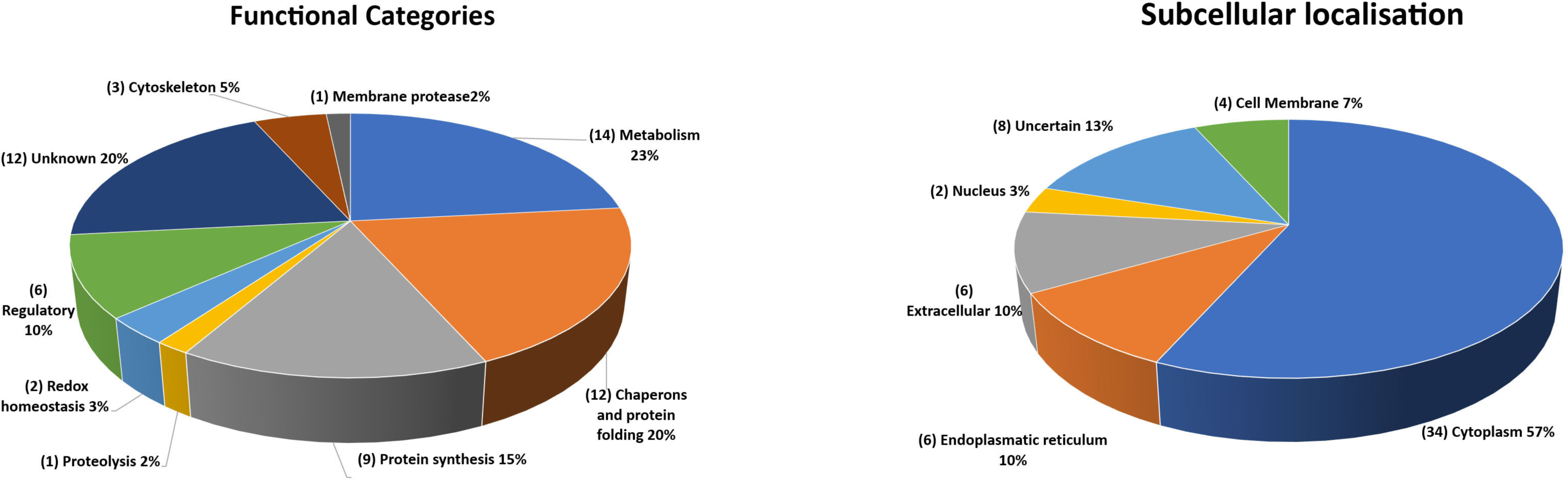
Quantitative representation of the different categories of the proteins identified in the extracellular vesicles. A, graphic of the EVs proteins distributed on the basis of their presumptive functions. The classification was based on the sequence homologies with characterized proteins. B, graphic of the EVs proteins distributed based on their presumptive sub-cellular localization. Prediction was made with DeepLoc-1.0 (http://www.cbs.dtu.dk/services/DeepLoc/).

All the proteins were also analysed in silico to identify signal peptide, transmembrane domain and endoplasmic reticulum retention signal, which are indicative traits of their subcellular localization. Thus, the largest group was composed of soluble components (34 in number) with a putative cytoplasmic localization (Figure 11, panel B). A second group of 7 proteins showed a signal for sorting and/or retention in the endoplasmic reticulum. Six secretory proteins were identified due to signal peptide at their N-terminus, whereas three showed a transmembrane domain. Finally, 2 proteins were predicted as nuclear components. However, the location of eight proteins remained unclear, as the potential location is assigned to subcellular compartments (*i.e.* mitochondrion, plastid, and lysosome), the existence of which is still unknown in *C. parvum*.

A putative interactome of EVs proteins was modelled by STRING interaction analysis. This search revealed a big network connecting 51 components (Supplemental figure S2, panel A and Supplemental Table S3), which was identified with a with medium confidence (0.4). The involvement of most (85%) of the identified proteins in this network emphasized the occurrence of a functional assembly bridging different molecular processes, which seemed highly represented in EVs. When higher confidence (0.9) was used for the analysis, this interaction network was limited to include only 34 proteins (57.3%) and three main sub-networks (Supplemental Figure S2, panel B) related to carbohydrate metabolism. Therefore, the above-reported interaction data highlighted the possible existence of functional macromolecular complexes in the *C. parvum* EVs involved in specific biological processes, as evidenced by the coherence with the most represented predicted functional categories (Figure 11, panel A).

The occurrence of highly related protein homologues assigned to EVs in other organisms was verified for 48 out of the 60 species identified in this study. Conversely, this proteomic investigation originally identified 12 proteins never assigned to EVs so far, among which all the ones (4 in number) here predicted being membrane components (Supplemental Table S2). An *in-silico* prediction of the schematic structure of these 12 proteins is shown in Figure 12. The occurrence of a signal peptide was evidenced in 10 proteins, among which 4 also contained a transmembrane domain, thus suggesting an extracellular/membrane localization for these components; the remaining 2 proteins did not contain localization motifs, as typical of cytoplasmic/luminal proteins. These 12 proteins included: i) three large molecules, such as the cysteine-rich secretory protein (cgd7_4310-RA-p1) including repeated CAP domains related to secretory proteins of metazoans (Gibbs et al., 2008) and allergen V5/Tpx-1-related (cgd5_2020_RA-p1); ii) a protein (cgd6_1630_RA) containing a CS domain, which is a binding module for HSP90, possibly involved in recruiting heat shock proteins to multiprotein assembly (Lee YT, et al 2004); iii) membrane-linked aspartyl-protease (cgd1_3690-RA-p1) that was detected only in the SEVs proteome. Additional proteins were uncharacterized proteins (cgd1_660-RA, cgd2_3780-RA, cgd3_1540-RA, cgd3_3370-RA, cgd6_1080-RA, cgd6_4460-RA, cgd6_710-RA, cgd7_400-RA) that do not show similarity with already known protein families. This group of *Cryptosporidium*-specific proteins also included the glycoprotein GP60 (cgd6_1080-RA-p1), which is a well-known immunodominant antigen of the parasite (Wanyiri et al., 2007) and was detected in both types of EVs.

**Figure 12.**
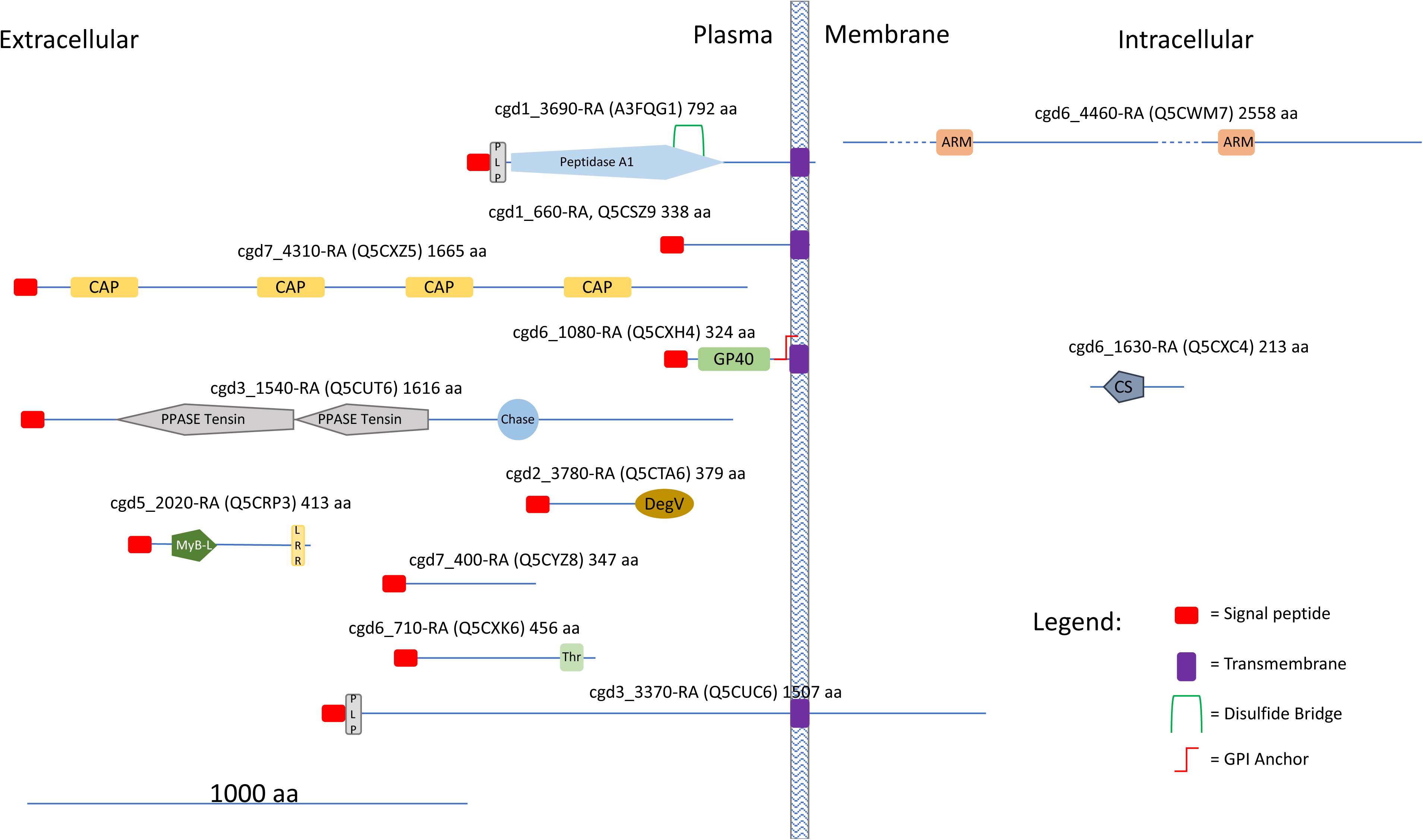
Cartoon showing some structural features of the *Cryptosporidium* characteristic proteins (see text). UniProtKB IDs are reported in brackets. Proteins were divided into three categories according to the presence of hydrophobic motifs (*i. e.* presence/absence of a signal peptide and/or a transmembrane domain) and their presumptive localization respect to the cell membrane. Prediction of signal peptides and transmembrane domains was performed at Phobius (https://phobius.sbc.su.se/index.html). Prediction of other structural and functional domains was performed at Pfam (http://pfam.xfam.org) and Prosite (https://prosite.expasy.org). List of domains in the figure: Peptidase 1, PEPTIDASE_A1, <u>PS51767</u>; PLP: PROKAR_LIPOPROTEIN, <u>PS51257</u>; CAP, Cysteine-rich secretory protein family, <u>CL0659</u>; GP60, Glycoprotein GP60 of Cryptosporidium, <u>PF11025</u>; PPASE Tensin, PPASE_TENSIN, <u>PS51181</u>; Chase, CHASE, <u>PS50839</u>; DegV, DEGV, <u>PS51482</u>; MyB-L, MYB_LIKE, <u>PS50090</u>; LRR, Leucin Rich Repeat, LRR, <u>PS51450</u>; Thr, Threonin Rich Region, THR_RICH, <u>PS50325</u>; ARM, Armadillo/plakoglobin ARM repeat, ARM_REPEAT, <u>PS50176</u>;CS, CS, <u>PS51203</u>.

## 4 Discussion

The excystation in *Cryptosporidium* spp. represents the beginning of a new infectious cycle and implies the release of intact vesicles from sporozoites. It has been previously demonstrated that micronemes and dense granules excrete the most of their soluble contents in a couple of hours after the excystation in the external medium (Chen et al., 2004). We here show that *C. parvum* sporozoites also release extracellular vesicles and that can be distinguished in large and small types, here named LEVs and SEVs.

Once the excystation starts, the release of EVs proceeds autonomously without any additional requirements, such as host factors, and can be readily achieved also *in vitro*. On this basis, differential centrifugation of the excystation medium resolved *C. parvum* EVs in two sediments that, based on their features, were related to LEVs and SEVs, respectively. Dedicated TEM experiments definitively demonstrated that these vesicles have a different size and, accordingly, can be distinguished into LEVs (medium size150 nm) and SEVs (medium size 60 nm). Furthermore, these vesicles are partially different for protein composition as initially shown by the western blotting analysis, which demonstrated the presence of CpRom1 and CpGRASP exclusively in LEVs, as well as by fluorescent labelling combined with SDS-PAGE analysis that revealed distinct protein banding.

Furthermore, the two types of vesicles are partially different in compositions as shown by the presence of CpRom1 and CpGRASP exclusively on LEVs as well as by the distinct fluorescent protein banding showed by the SDS-PAGE analysis. This partial difference was finally verified by dedicated proteomic experiments that traced some proteins only in one of the two sediments; in particular, 39 proteins were in common between two vesicle types, while 5 and 16 proteins were identified only in LEVs and SEVs, respectively.

The finding of *C. parvum* EVs reported in the present study was fortuitous, and it was the consequence of our investigation on CpRom1 rhomboid, which belongs to a peculiar family of serine proteases integrally embedded in the plasma membrane. Apicomplexa have different types of rhomboids that act cleaving specific membrane proteins in a sequential manner; some of these molecules have fundamental roles in the motility of the parasite, and in its penetration capacity into the host cell (Dowse et al., 2008; Rugarabamu et al., 2015). At present, the specific role of CpRom1 in sporozoite invasion of enterocytes is still unclear. Nevertheless, this study confirmed that mature CpRom1is the largest rhomboid described so far, with a measured mass of 110 kDa. The two-dimensional model of this protein predicted a distinctive feature, with a large N-terminal hydrophilic portion extending outward from the plasma membrane (Figure 2). Consistent with this model, the gold particles used in immune electron microscopy experiments labelled the external membrane of LEVs (Figure 4B). These results were obtained using antibodies targeting the ID2 antigen, which is part of the above-reported N-terminal region exposed on the external surface of the membrane (Figure 2). It is worth noting that, except for those with mitochondrial localization (PARLs), all the rhomboids described so far are associated with the external side of cell membrane (Kandel and Neal, 2020). In this context, this study is original since it demonstrates for the first time the association of a rhomboid with extracellular vesicles.

To determine the subcellular origin of the sporozoite EVs, we sought to trace proteins that are undoubtedly involved in vesicular trafficking within the cell. Among these proteins, we observed that the GRASP proteins from *Cryptosporidium* spp. are strictly related to other members of this protein family. Indeed, immunoblotting experiments demonstrated that CpGRASP, a member of the conserved Golgi-associated protein family, is present on LEVs and not on SEVs similarly to CpRom1. Golgi reassembly stacking protein family are tethered to the external membrane of Golgi vesicles, forming the Golgi stack network that coordinates the sorting of secreted proteins (Rabouille and Lindsted, 2016). In addition, GRASPs regulate the unconventional secretion of proteins directed toward the cell membrane, tagging vesicles originating directly from the ER through a Golgi-independent pathway (Ahat et al., 2019). Therefore, the exclusive presence of CpGRASP on LEVs suggests that the biogenesis of these vesicles is associated with the ER- and/or the Golgi pathway.

TEM and SEM micrographs clearly showed the generation of EVs budding from the external membrane of sporozoites (Figure 7). Budding from the plasma membrane is the most common mechanism for the generation of EVs, except for exosomes. Among other things, a specific difference between budding-generated EVs (or ectosomes) and exomes is their size, which ranges from 100 to 1000 nm for the ectosomes and from 30 to 150 nm for the exosomes (Meldolesi, 2018). Indeed, the newly formed vesicles as shown in Figure 7B and 7D were larger than 150 nm, in agreement with what expected for vesicles generated through the budding process. Therefore, we can conclude that outward budding of *C. parvum* sporozoite membrane is responsible for the generation of LEVs.

Tentatively determining the origin of smaller SEVs was more challenging, as these vesicles were only observed after their release into the excystation medium. Since SEVs share some physical parameters with exosome like density (ranging from 1.08 to 1.2 g/ml) and size (ranging from 50 to 150 nm) it was reasonable to consider SEVs as exosome-like vesicles. Exosomes are generally distinguished from other vesicles for their origin within the endosomal pathway through the formation of MVBs; upon fusion with the plasma membrane these intracellular bodies release exosomes outside the cell (Doyle and Wang, 2019). After assuming the existence of organelles like MVBs in sporozoites, we accurately examined the corresponding TEM micrographs finally identifying novel cytoplasmic structures that highly resemble MVBs; accordingly, we here defined them MVB-like organelles. Analogously to MVBs, MVB-like organelles are made of a large globular vesicle enclosed by a lipid membrane, which in turn contains various smaller vesicles (Figure 8). These novel organelles occur in proximity of the sporozoite membrane. However, since MVBs of proven endosomal origin have not yet been described in *Cryptosporidium* spp., our assimilation of SEVs to exosomes remains speculative.

Integrated Western blotting, fluorescent labelling and proteomics experiments have shown that *C. parvum* LEVs and SEVs also differ slightly in protein composition. Of the 60 proteins assigned to proteomics, 39 were common in two types of vesicles, while 5 and 16 components were unique to LEVs and SEVs respectively. However, proteins identified by proteomic analysis represent the most abundant proteins of the vesicles and other methods (*i.e.* Western blotting) can be used to track lower-enriched proteins such as CpRom1 and CpGRASP that selectively occur in LEVs.

Based on a specific analysis for presumptive cellular localization, most (35 out of 60) of these vesicle proteins were predicted to occur in the cell cytoplasm. Various types of cytoplasmic proteins have already been described in extracellular vesicles (Gurung et al., 2021), even if their function in these particles is not yet clear. However, it is conceivable that some cytoplasmic proteins are dragged from the cytoplasm during the formation of the emerging vesicles. On the other hand, 6 vesicle proteins here identified show a consensus sequence for the endoplasmic reticulum (ER), while other 6 ones contain a signal peptide, suggesting their sorting towards EVs through specific ER and/or Golgi-mediated secretory pathways.

Based on gene ontology information, most (48 out of 60) of the proteins here identified were tentatively assigned to a known functional category, while 12 ones were considered peculiar of *Cryptosporidium* spp. Among the former group, the largest number of annotated molecules included enzymes related to cellular metabolism and components involved in protein folding and biosynthesis.

Bioinformatic analysis of the corresponding protein interactions revealed the existence of a single network connecting most (51 out of 60) of the assigned molecular entries, suggesting the existence of unique functional machinery interconnecting different processes.

Regarding the remaining 12 *Cryptosporidium*-specific vesicle proteins, 10 have never been described so far, whereas the protein identified as cgd1_3690-RA was clearly homologous to an aspartyl-protease and glycoprotein GP60 (cgd6_1080-RA) have been already mentioned in several studies. Figure 12 outlines some of the molecular features of these *Cryptosporidium*-specific proteins, including their presumptive localization with respect to plasma membrane, which is based on the presence of a signal peptide and transmembrane or other specific domains. Noteworthy, 10 of these proteins show a signal peptide, suggesting that reach the vesicle through a secretive pathway, while 4 of them, such as GP60, have a transmembrane domain essential for their exposure on the surface of EVs.

With respect to aspartyl protease (cgd1_3690-RA), which we will call membrane protease CpMAP, it has only been detected in SEVs. *C. parvum* aspartyl-proteases were previously demonstrated being crucial for a successful infection. In this context, the aspartyl-protease inhibitor Indinavir was previously shown reducing parasite proliferation both *in vitro* and *in vivo* when administered in the early phase of infection (Mele et al., 2003). The present study demonstrates that CpMAP is expressed in the early stage of parasite infection, and it is conveyed by vesicles outside the sporozoites, thus possibly playing a role in the proteolysis of one or more host proteins.

Otherwise, GP60 was detected in LEVs and SEVs. This glycoprotein, also referred to as GP40/15, is one of the most relevant antigens of the parasite, which may determine an immunodominant response in the host (Sestak et al., 2002). It occurs exclusively in the *Cryptosporidium* genus and has no homologs even in other apicomplexans. This protein is synthesized as a large precursor of 60 kDa (GP60), which may undergo a proteolytic cleavage by a furin-like protease to generate a soluble fragment of approximately 40 kDa (GP40) and a smaller membrane-anchored segment of 15 kDa (GP15) (Wanyiri et al., 2007). Despite its nature of membrane protein, GP15 is released during the sporozoite gliding (O’Connor et al., 2007). It is remarkable that this fact is in perfect accordance with the discharge of EVs after the excystation.

Extracellular vesicles constitute important elements of the host-parasite interaction and mediate cell-to-cell communication in different directions: between the parasite and its host, among the parasites, and among the host cells in response to the parasitic infection (Wu et al., 2019). In most of the cases, parasite-generated vesicles modulate the host immune response by transferring parasite molecules (*i.e.* mRNA, various types of non-coding RNA, DNA and proteins) that act on host cells like macrophages (Olajide and Cai, 2020).

In apicomplexan parasites, extracellular vesicles were studied in *Plasmodium* spp. and *Toxoplasma gondii*. In *Plasmodium* spp. host-generated exosomes have been described containing parasite proteins. In *Plasmodium yoelii*, these exosomes inoculated in mice can elicit an IgG response and a protective immunity (Martin-Jaular et al., 2011). In *Plasmodium falciparum*, host-generated exosomes can also coordinate gametocytogenesis among infected red blood cells by delivering parasitic DNA (Regev-Rudzki et al., 2013). Importantly, microvesicles induced by *Plasmodium* spp. contribute to the onset of inflammatory symptoms of malaria. In *P. falciparum,* the number of extracellular vesicles excreted by blood cells increases in cerebral malaria (Pankoui Mfonkeu et al., 2010; Sahu et al., 2013), and a high number of them correlates with high fever in *Plasmodium vivax* infection (Campos et al., 2010).

Host-generated extracellular vesicles are induced also by *T. gondii* in infected fibroblasts (Pope and Lässer, 2013), and the parasitic antigens in microvesicles stimulate dendritic cells leading to a Th1 protective response (Dlugońska and Gatkowska, 2016). It was also demonstrated that *T. gondii* generates its own exosomes and extracellular vesicles (Wowk PF et al., 2017).

With this study, we have added novel information in the complex scenario of extracellular vesicles secreted by apicomplexan parasites. We discovered that also *C. parvum* excysted sporozoites release EVs. Based on their size, these extracellular vesicles were resolved in LEVs and SEVs. Integrated approaches allowed defining common and peculiar characteristics of both vesicle typologies. CpRom1 and CpGRASP were selectively identified in LEVs, and thus can be considered markers of this vesicle type. Conversely, the abundant antigenic glycoprotein GP60 was detected in both type of vesicles; this fact may play a role in directing the host’s immune response. Further studies are needed to decipher the specific function of these extracellular vesicles and their relationship with *C. parvum* infection.

## Author contributions

LB: Investigation, Methodology, Writing – review & editing. ZB: Investigation, Methodology, Writing – review & editing. AMS: Investigation, Methodology, Writing – review & editing. IV: Investigation, Methodology, Writing – review & editing. IP: Investigation, Writing – review & editing. EM: Investigation, Writing – review & editing. AS: Conceptualization, Formal Analysis, Writing – review & editing. MS^1^: Conceptualization, Formal Analysis, Writing – review & editing. MS^6^: Conceptualization, Formal Analysis, Writing – review & editing. MLF: Supervision, Conceptualization, Formal Analysis, Writing – review & editing. FT: Resources, Supervision, Conceptualization, Investigation, Methodology, Writing – review & editing.

## Funding

This work was supported by grants from the National Recovery and Resilience Plan, mission 4, component 2, investment 1.3, MUR call n. 341/2022 funded by the European Union-NextGenerationEU for the project “One Health Basic and Translational Research Actions addressing Unmet Needs on Emerging Infectious Diseases (IN-FACT)”, D.C. 1554/2022, PE00000007.

## Acknowledgments

We are thankful to CryptoDB for making available all the genomic data used in this study.

## Conflict of Interest

The authors declare that the research was conducted in the absence of any commercial or financial relationships that could be construed as a potential conflict of interest.

## Notes

### Competing Interest Statement

The authors have declared no competing interest.

### Summary of Updates

To correct the order of figures in the manuscript.

