## Supplementary figures and images for "Unveiling *Cryptosporidium parvum* Sporozoite-Derived Extracellular Vesicles: Profiling, Origin, and Protein Composition"

### Supplemental Figure S1

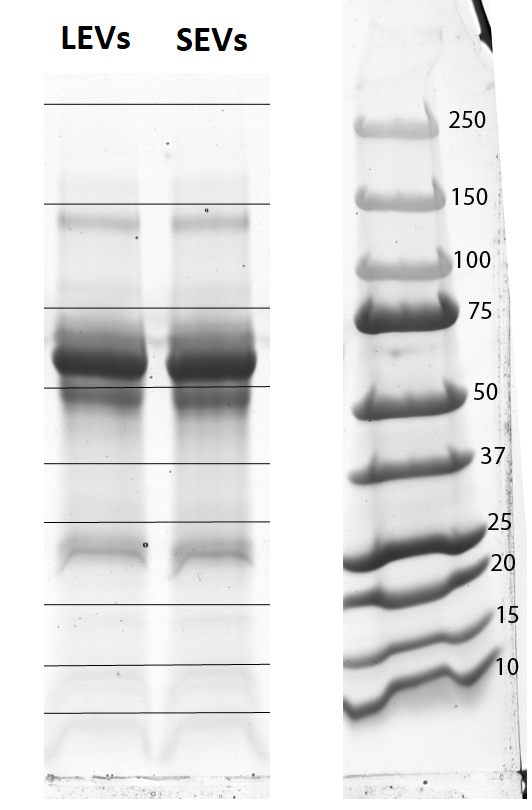

### Supplemental Figure S2

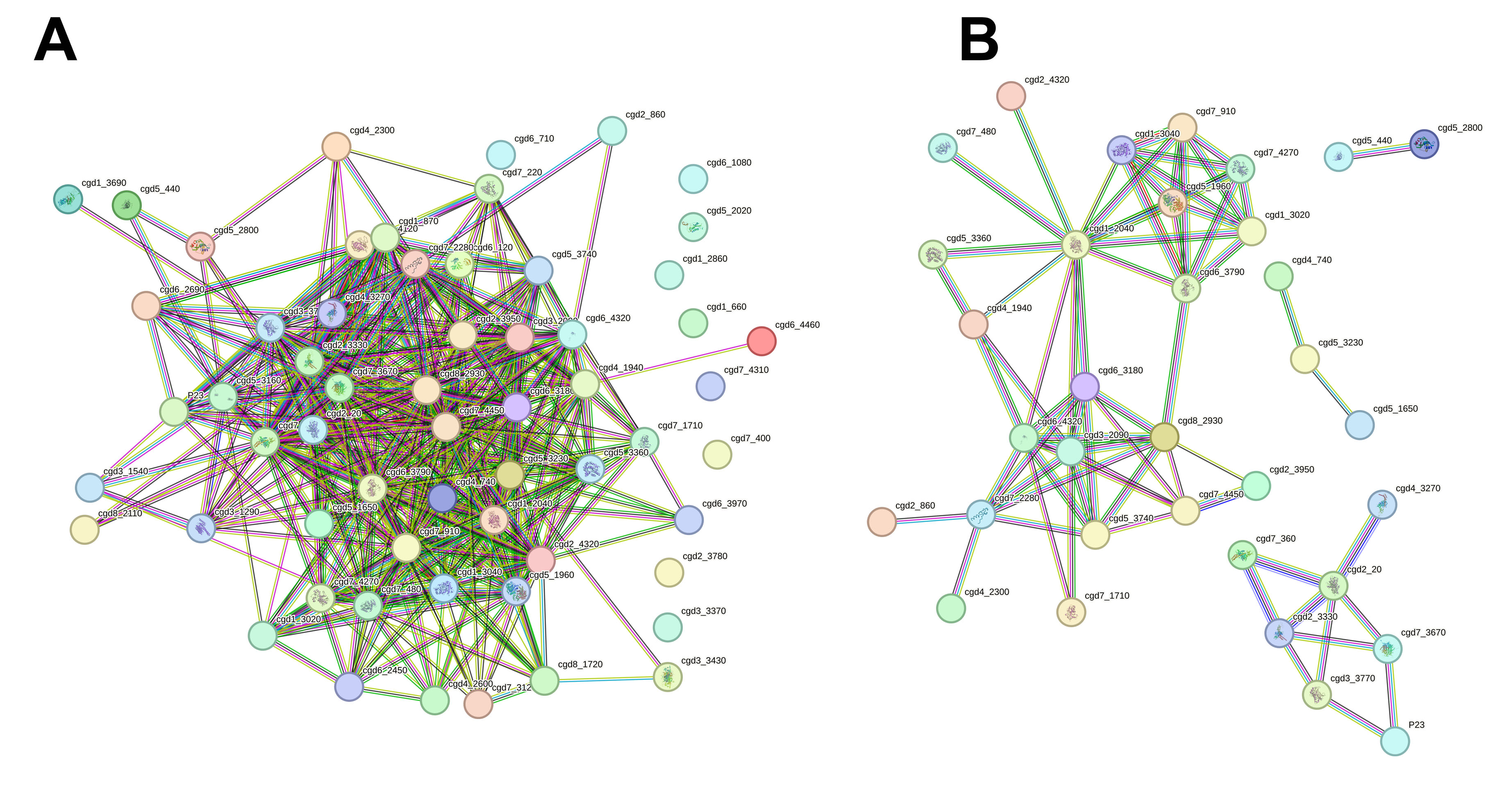
