## Supplemental Table S3 for "Unveiling *Cryptosporidium parvum* Sporozoite-Derived Extracellular Vesicles: Profiling, Origin, and Protein Composition"

| **Accessione code**  **CryptoDB** | **String Name** | **Annotation** |
| --- | --- | --- |
| cgd7_480-RA-p1 | Q5CYZ2 | Lactate dehydrogenase, adjacent gene encodes predicted malate dehydrogenase |
| cgd4_2600-RA-p1 | A3FQL3 | UDP-glucose 4-epimerase |
| cgd1_660-RA-p1 | Q5CSZ9 | Predicted secreted protein, signal peptide |
| cgd3_1540-RA-p1 | Q5CUT6 | Large protein with signal peptide. cysteine-rich, threonine-rich, possible mucin |
| cgd6_4460-RA-p1 | Q5CWM7 | Large protein with ARM repeats |
| cgd1_3040-RA-p1 | Q5CSE7 | Triosephosphate isomerase |
| cgd5_3740-RA-p1 | Q5CQ87 | 40S ribosomal protein S12 |
| cgd2_4320-RA-p1 | Q5CT58 | Thioredoxin reductase 1 |
| cgd4_3270-RA-p1 | A3FQM0 | Heat shock 105kD heat shock 105kD alpha heat shock 105kD beta heat shock 105kDa protein 1 |
| cgd4_740-RA-p1 | A3FQN6 | Thioredoxin peroxidase-like protein |
| cgd1_870-RA-p1 | Q5CSY2 | Peptidyl-prolyl cis-trans isomerase |
| cgd7_400-RA-p1 | Q5CYZ8 | Uncharacterized protein |
| cgd7_1710-RA-p1 | Q5CYN0 | Threonyl-tRNA synthetase (RNA binding domain TGS+HxxxH+tRNA synthetase) |
| cgd7_910-RA-p1 | Q5CYV1 | Phosphoglycerate kinase |
| cgd5_3360-RA-p1 | Q5CRC5 | Adenylate kinase |
| cgd6_3790-RA-p1 | Q5CWT6 | Glyceraldehyde-3-phosphate dehydrogenase |
| cgd5_1960-RA-p1 | Q5CRP8 | Enolase (2-phosphoglycerate dehydratase) |
| cgd7_4310-RA-p1 | Q5CXZ5 | Extracellular protein with a signal peptide and 4 SCP domains |
| cgd3_2090-RA-p1 | Q5CUN6 | 40S ribosomal protein SAe |
| cgd2_3330-RA-p1 | Q5CTE6 | APG-1 like HSP70 domain containing protein, signal peptide plus likely ER retention motif |
| cgd4_2300-RA-p1 | Q5CR33 | Ubiquitin-activating enzyme E1 (UBA) |
| cgd8_1720-RA-p1 | Q5CW41 | Acetaldehyde reductase plus alcohol dehydrogenase (AdhE) of possible bacterial origin |
| cgd1_2040-RA-p1 | Q5CSM7 | Pyruvate kinase |
| cgd5_3160-RA-p1 | A3FQJ8 | Actin |
| cgd8_2930-RA-p1 | Q5CVS6 | Eft2p GTpase translation elongation factor 2 (EF-2) |
| cgd6_1080-RA-p1 | Q5CXH4 | GP40 domain-containing protein |
| cgd3_1290-RA-p1 | Q5CUW0 | 14-3-3 domain containing protein |
| cgd3_3430-RA-p1 | Q5CUC0 | Amine oxidase |
| cgd4_1940-RA-p1 | Q5CR64 | Putative nucleoside-diphosphate kinase |
| cgd7_3670-RA-p1 | Q5CY50 | Heat shock protein 90 (Hsp90), signal peptide plus ER retention motif |
| cgd1_3020-RA-p1 | Q5CSE9 | Fructose-bisphosphate aldolase |
| cgd5_2020-RA-p1 | Q5CRP3 | Extracellular protein |
| cgd6_2450-RA-p1 | Q5CX54 | Alpha-1,4 glucan phosphorylase |
| cgd7_2280-RA-p1 | Q5CYH8 | 60S ribosomal protein L40 |
| cgd6_120-RA-p1 | Q5CPJ7 | Disulfide-isomerase, signal peptide plus ER retention motif, putative ER protein |
| cgd7_4270-RA-p1 | Q5CXZ9 | Phosphoglycerate mutase |
| cgd7_4450-RA-p1 | Q5CXY4 | Elongation factor EF1-gamma (Glutathione S-transferase family) |
| cgd6_2690-RA-p1 | Q5CX33 | Peptidylprolyl isomerase |
| cgd2_20-RA-p1 | Q5CPP8 | Heat shock 70 (HSP70) protein |
| cgd5_2800-RA-p1 | Q5CRH0 | Actin depolymerizing factor |
| cgd3_3370-RA-p1 | Q5CUC6 | Uncharacterized protein |
| cgd3_3770-RA-p1 | Hsp90 | Hsp90 |
| cgd6_710-RA-p1 | Q5CXK6 | Protein with signal peptide plus Thr stretch, possible mucin |
| cgd2_3950-RA-p1 | Q5CT92 | Putative translation elongation factor 1 beta 1 |
| cgd6_1630-RA-p1 | P23 | P23 HSP20-like chaperones fold |
| cgd8_2110-RA-p1 | Q5CW04 | Uncharacterized protein |
| cgd2_4120-RA-p1 | A3FQA7 | Peptidyl-prolyl cis-trans isomerase |
| cgd6_3180-RA-p1 | Q5CWZ0 | 40S ribosomal protein S15 |
| cgd6_4320-RA-p1 | Q5CWP0 | 40S ribosomal protein S5 |
| cgd7_360-RA-p1 | Q5CZ02 | Heat shock protein, Hsp70 |
| cgd5_1650-RA-p1 | A3FQI3 | DJ-1_PfpI domain-containing protein |
| cgd1_3690-RA-p1 | A3FQG1 | Aspartyl (Acid) protease, putative |
| cgd6_3970-RA-p1 | Q5CWS1 | Glutaredoxin-like protein 2 thioredoxin folds |
| cgd1_2860-RA-p1 | A3FQF3 | Uncharacterized protein |
| cgd5_3230-RA-p1 | A3FQJ9 | Superoxide dismutase |
| cgd7_220-RA-p1 | A3FPL8 | GTP-binding nuclear protein |
| cgd5_440-RA-p1 | Q5CS32 | Protein with 2 CAP (CARP) domains, possible adenyl cyclase-associated protein |
| cgd7_3120-RA-p1 | Q5CYA3 | Pyruvate decarboxylase |
| cgd2_3780-RA-p1 | Q5CTA6 | Uncharacterized protein rich in Thr and Ser residues |
| cgd2_860-RA-p1 | Q5CU00 | Proteasome subunit beta |
